# Protein homeostasis imprinting across evolution

**DOI:** 10.1101/2020.07.19.210591

**Authors:** Thodoris Koutsandreas, Brice Felden, Eric Chevet, Aristotelis Chatziioannou

**Author notes:** equally contributed to this work (Brice Felden, Eric Chevet, Aristotelis Chatziioannou). in memoriam of Prof. Brice Felden. ***Corresponding Authors:*** TK, EC, AC.

## Abstract

Protein homeostasis (a.k.a. proteostasis) is associated with the primary functions of life, and therefore with evolution. However, it is unclear how the cellular proteostasis machines have evolved to adjust the protein biogenesis needs to environmental constraints. Herein, we describe a novel computational approach, based on semantic network analysis, to evaluate proteostasis differentiation during evolution. We show that the molecular components of the proteostasis network (PN) are reliable metrics to deconvolute the life forms into Archaea, Bacteria and Eukarya and to assess the evolution rates among species. Topological properties of semantic graphs were used as new criteria to evaluate PN complexity of 93 Eukarya, 250 Bacteria and 62 Archaea, thus representing a novel strategy for taxonomic classification. This functional analysis provides information about species divergence and pointed towards taxonomic clades that evolved faster than others. Kingdom-specific PN were identified, suggesting that PN complexity correlates evolution. Through the analysis of gene conservation, we found that the gains or losses that occurred throughout PN evolution revealed a dichotomy within both the PN conserved modules and within kingdom-specific modules. Since the PN is implicated in cell fitness, aging and disease onset, it could be used as a new metric to tackle mechanisms underlying ‘gain-of-functions’, and their biological ramifications.

## Introduction

Protein homeostasis (a.k.a. proteostasis) refers to a complex and interconnected network of processes that affects both expression levels and conformational stability of proteins, by controlling their biogenesis, folding, trafficking and degradation within and outside the cell. The molecular mechanisms controlling proteostasis are implicated in cell fitness, aging and contribute to disease onset. The underlying network of cellular mechanisms (i.e. the proteostasis network - PN) includes protein synthesis, co/post-translational folding, quality control, degradation, as well as adaptive signaling in response to proteostasis imbalance (Balch et al., 2008). From Prokaryotes to Eukaryotes, the PN was subjected to evolutionary pressure for each organism, to cope with intrinsic and extrinsic demands. Evolution was shaped by factors such as genome complexity, post-translational modifications repertoire, the presence of subcellular compartments, the emergence of multicellular organisms and cell differentiation, with cells exhibiting high protein synthesis yields and secretion, requiring a robust PN. An effective PN is also instrumental for eukaryotic cells with temporal variations of protein expression (e.g. neurons and endocrine cells). Each of these constraints increased the needs for updated, adaptive mechanisms, ensuring protein homeostasis (Roth & Balch, 2011). As such, the PN was subjected to selections, specific to each organism. For instance, compartment-specific proteostasis control machineries in the cytosol, the endoplasmic reticulum (ER) or the mitochondria are required in eukaryotic cells, whereas dedicated systems are found in the cytosol and periplasm of Gram-negative Bacteria (Powers & Balch, 2013). Despite the fact that the proteostasis network is a crucial, biological module, with direct relevance to many diseases linked with aberrant protein conformation, yet its fragmented, problematic annotation, hinders the efficient investigation of its functional ramifications in health and disease (The Proteostasis Consortium, Overall coordination et al., 2022).

Traditional, phylogenetic approaches use the sequence of conserved genes or proteins (or groups of them) as standard references, rather than their functional identities, to form clades, ancestral lineages and identify speciation events. A proper PN-related phylogenetic clustergram needs to reflect the diversity of PN across species of various taxa. In theory, heat shock proteins (HSPs), that exert fundamental roles in maintaining protein homeostasis, could be considered as the appropriate markers to delineate PN evolution. Some HSP families (e.g. HSP40 and HSP70) are highly represented in most cells and the nature of this representation might reflect the underlying evolutionary relationships - e.g. 3 members of HSP40 in *E. coli* and 49 members in *Homo sapiens* (Gur et al., 2005; Kampinga et al., 2009). However, the content of their nucleotide and amino acid sequences remains unable to provide insights about the functional evolution of proteostasis. Hence, a different vocabulary is necessary to exploit their functional profiles and the subsequent structured networks.

Even if studies have reported computational models of proteostasis in *E. coli* Powers et al., 2012) and in Eukaryotes (Wiseman et al., 2007), an overall layout of proteostasis evolution is still lacking. Herein, we propose a novel method for phylogenetic analysis, based on the semantic network analysis of ontological graphs, which describes the functional characteristics of species, under a specific cellular process, such as proteostasis. Using this approach to evaluate the functions of specific biological modules (i.e. list of genes/proteins), we assessed how proteostasis evolved with the phylogeny. Proteostasis-based evolutionary maps performed well *vs*. the gold-standard ribosomal RNA (rRNA) sequence-based evolution metric (Gilbert, 1986; Smit et al., 2007). Specifically, these maps managed to yield a phylogenetic clustergram, which proposes taxonomic classifications of organisms, based on the topology of their PN. In addition, these newly generated clustergrams reproduce the established classifications in major taxa. Species allocation into distinct clusters, populated by members from various phyla, unveiled new shared, adaptive responses to environmental cues. We show here that proteostasis, as a whole, represents a reliable metric for species partitioning, providing a snapshot of the overall, functional diversification of cellular functionality, in the tree of life. Conserved and ‘kingdom-specific’ PN components were identified, implying that proteostasis is associated with a plethora of crucial, cellular functions, in different taxonomies, apart from its core machinery.

## Materials & Methods

### Proteostasis-related genes lists and data acquisition

The PN encompasses various mechanisms, pertinent to different functional aspects, as protein quality control, production, concentration maintenance and degradation. Moreover, proteostasis regulates a multitude of cellular processes and consequently has a powerful contribution to the large phenotypic diversity observed. This large complexity is partly documented by the available vocabularies of biological pathways and processes (e.g. Gene Ontology (The Gene Ontology Consortium, 2019), Reactome (Jassal et al., 2020), KEGG (Kanehisa & Goto, 2000)). Therefore, the semantic representation and annotation of the PN is largely skewed. On account of this, we defined species-specific gene lists, related to proteostasis, in order to reveal the PN components, utilizing the pathway analysis. This gene selection procedure was performed through a supervised, multi-phase, analytic workflow (**Fig. 1A, step I**), and it aimed to include genes strongly associated with key functional components, such as protein folding, degradation, endoplasmic reticulum, autophagy and associated signaling pathways. In this way, seed gene lists were defined for eight eukaryotic model species (*Homo sapiens, Gallus gallus, Danio rerio, Xenopus tropicalis, Caenorhabditis elegans, Drosophila melanogaster, Saccharomyces cerevisiae and Arabidopsis Thaliana*) and a generic gene list was created for the prokaryotic domain (see Code and Data Availability). For Eukaryotes, we applied homology mappings so as to retrieve putative, functionally similar genes, from the Ensembl database (vertebrates, fungi, metazoans and plants, Kinsella et al., 2011). The aforementioned model species were used to detect homologies, with species belonging to the same taxonomic classification level (e.g. *S. cerevisiae* was used as a reference species for fungi and *A. thaliana* for plants). Hence, representative proteostasis-related gene lists were constructed automatically for hundreds of Eukaryotes. However, in order to exclude spurious annotations, only species pertaining a number of genes, above 75% of the reference gene sets, were included into the final set.

**Figure 1:**
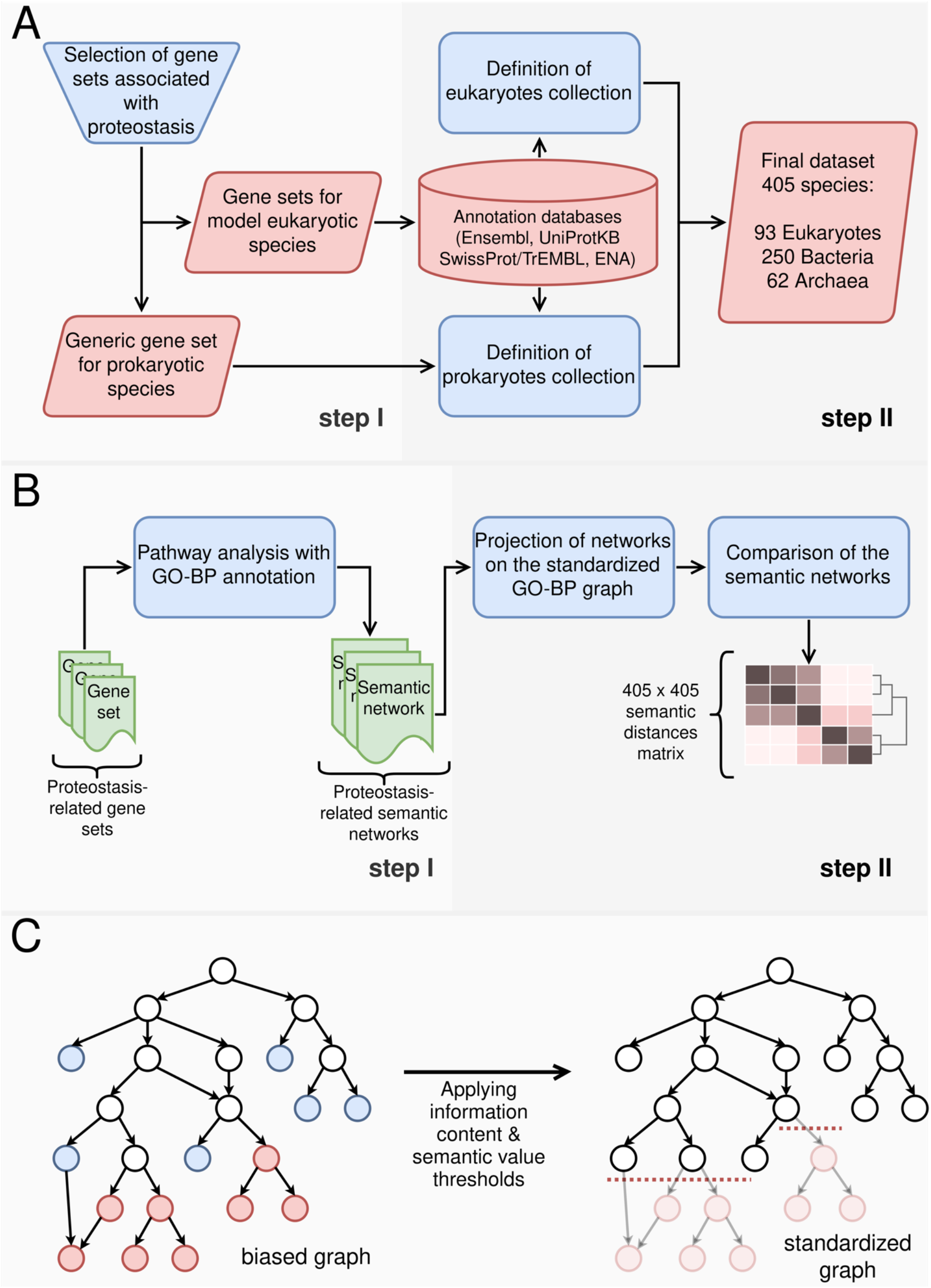
Schematic analytical workflow. **(A)** Process of data acquisition. Proteostasis-related gene lists were constructed manually for model organisms (step I). Then, public databases were used to collect genomic homologies to expand species collection and concentrate specific data for rRNA, HSP40 and HSP70 sequences. Only species with ample annotation were selected in the final set (step II). **(B)** Analysis workflow. Gene lists were translated into semantic networks through pathway analysis of GO-BP (step I). A standardized version of the GO-BP graph was constructed to remove potential annotation bias from the results. Finally, comparative analysis was performed to calculate the differences among proteostasis-related networks (step II). **(C)** Construction of the standardized GO-BP graph: Terms residing low in the branches of the graph bear high IC (red and blue nodes). Some of them are located in deep, densely populated branches (red nodes), reflecting the fact that these processes are more extensively studied than others. This imbalance in knowledge representation is a source of annotation bias, which was neutralized by filtering out terms in voluminous ontological regions (elongated and abundant branches), with high information content (red nodes).

The advantage of the genomic annotation of Prokaryotes, compared to that of Eukaryotes, is the adoption of a common nomenclature. Homologous genes in different species are referred with the same gene symbol, facilitating their automated search and association. Hence, the initial prokaryotic gene list was used as the basis to construct a proteostasis profile for thousands of species. These profiles were extracted from the UniProt Knowledgebase (The UniProt Consortium, 2018), which contains numerous reference proteomes of fully sequenced species. To focus on taxonomically proximal species with ‘good quality’ genomic annotations, only species with complete proteome detector (CPD index equal to “Standard”) were selected. As thousands of Bacteria met that criterion, a random selection was then performed to reduce their number to 250. The selection procedure adopted the constraint to select at least one member for each taxonomic Class. The taxonomic analysis, using as metric the topological properties of the PN, was contrasted to classical, taxonomic analyses, which utilize either the ribosomal sequences (18S and 16S rRNAs), retrieved from the ENA repository (Leinonen et al., 2010), or heat shock proteins data (HSP40 and HSP70), collected from Ensembl and UniProt Knowledgebase (**Fig. 1A, step II**).

### Pathway analysis of proteostasis-related genes lists

A PN semantic profile was constructed for each species, based on the Biological Process corpus of Gene Ontology (GO-BP) **(Fig. 1B, step I**). We used the prioritized pathway analysis described in Koutsandreas et al., 2016, while the whole implementation can be found in a publicly available Galaxy-engine platform (http://www.biotranslator.gr:8080/). The algorithm utilizes the statistical distribution of enrichment scores, derived from the respective gene lists of each species, which is reordered according to the frequency of the scores, further corrected by non-parametric, permutation resampling so as to prioritize the final set of enriched pathways. For each species, this semantic processing identified a network of biological processes, delineating the semantic tree of proteostasis. We used two criteria to determine the enriched biological processes. Hypergeometric p-value cut-off was set to 0.05 and then, the GO-BP terms were prioritized according to the adjusted p-value. The cut-off for the adjusted p-value was set at 0.05, however if a species had fewer than one hundred terms satisfying that threshold, the selection was extended to the first hundred terms to keep a comparable cardinality, among the sizes of lists.

### Standardized GO-BP and PN profiles

The GO-BP graph represents a vocabulary of biological knowledge, which is used to provide a genome-wide interpretation of experimental results. Yet, inherent inconsistencies regarding the structure and the depth of its branches generate bias that hampers comparative analysis, between different species (Gaudet & Dessimoz, 2017). Some graph branches are more expanded than others, due to extensive annotation. This leads to distorted, descriptive capacity, regarding the degree of semantic specification that each term bears. Namely, graph branches do not carry the same, semantic weight. Furthermore, the depth of genomic annotation differs among species. Different research communities have developed rich, genomic annotations for model species, emphasizing on specific components of cellular physiology, according to specific, operational or developmental characteristics of each organism (Gaudet & Dessimoz, 2017). On the other hand, the vast majority of species has been sketchily annotated, only through electronic inference, based on homology associations (Gaudet et al., 2011). This causes inconsistencies regarding the semantic network profile across species, and even taxonomically related species, could have divergence in annotation coverage (Lobb et al., 2020). All these predict upon a standardized version of GO, suitable for comparative analyses among various species. The construction of such a standardized, unbiased graph relies on two metrics of ontological graphs, information content (IC) and semantic value (SV). IC measures the specificity of a term, considering the amount of its descendant nodes (Pesquita et al., 2009). Conceptually, a term with plenty of descendant nodes has low IC value because its semantic content can be further broken down into more specific concepts. On the other hand, graph leaves have the maximum IC value. IC is defined as follows:

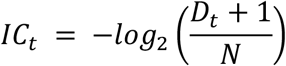

where *D_t_* refers to the number of descendants of term *t* and *N* is the cardinality of the complete set of terms. As it considers only the number of its child-terms, IC does not integrate the topological information of a term. For example, all leaf nodes have the same IC, but are located at different depths within the graph, depending on the quality of annotation. SV is a metric, proposed to overcome that limitation. It depicts the semantic distance of a term from the root, considering the information contained in its ancestral plexus (X. Song et al., 2014):

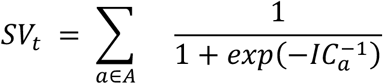

where *A* is the set of ancestors of term *t*. The SV of a term is linearly correlated to the amount of its ancestors. High values point out either increased distances from the root or the existence of extensively described semantic regions. The latter causes annotation bias favoring the overpopulated tree branches.

For the purposes of the comparative analysis, we created a semantic graph that would render feasible the comparison among species, by trading off between the detail of annotation of semantic branches and the need to minimize annotation bias. We constructed a standardized version of GO-BP, by filtering out very specific terms (high IC) from expanded ontological areas (high SV). Terms exceeding the twentieth (20^th^) percentiles of IC and SV distributions were trimmed and substituted with their most proximal ancestors, conforming to these thresholds (**Fig. 1C**). The initial set of 20596 annotated GO-BP terms was decreased to 2693 terms.

The PN profiles derived from pathway analysis were transformed within the terms of this standardized GO-BP graph. Specifically, enriched terms not included in the corpus of the standardized graph were mapped to ancestral terms that existed in the graph. For example, the term “peptidyl-proline modification” (GO:0018208) was substituted with “protein metabolic process” (GO:0019538) and “macromolecule modification” (GO:0043412). On the other hand, each enriched term included in the standardized graph was mapped to a list, which contained only itself. As a result, each PN profile was transformed from a list of enriched terms into a set of term lists. These new semantic representations were used for the comparative analysis, as well the identification of PN components (**Fig. 1B, step II**).

### Comparative analysis of the PN profiles

The calculation of the group-wise semantic similarities of the PN profiles was based on the exploitation of GO-BP structure. Initially, terms with information content lower than 0.1 were filtered out from the enriched sets. These terms referred to very generic biological processes (e.g. biological process, biological regulation, etc) and their inclusion in the PN profiles suppresses the potential semantic difference between two species. All other enriched terms were mapped to their respective sets of terms included in the standardized graph, as described in the previous section.

In general, semantic comparison estimates the closeness of two ontological terms, and is based on the topological relevance of their ancestors (pairwise measures). Due to the mapping of the enriched terms to sets of terms in the standardized graph, we adjusted the concept of semantic comparison to be applicable on groups of terms. Initially, a global similarity matrix was constructed for all the terms of standardized GO-BP. To avoid bias of specific pairwise measures, the similarity of two terms was calculated by averaging three widely used metrics: Resnik (Resnik, 1999), XGraSm (Mazandu et al., 2016) and AggregateIC (X. Song et al., 2014). Then, the similarity of two enriched terms was calculated as the mean similarity of the respective sets of terms in the standardized graph. In this way, we estimated the pairwise similarities of enriched terms. Finally, the similarity of two species was calculated based on the average best matches formula (Mazandu et al., 2016):

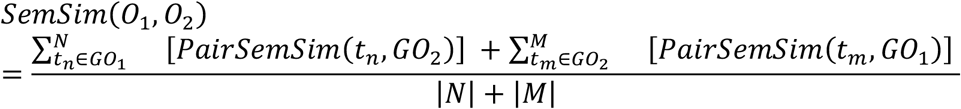

where *G*_1_ and *GO*_2_ are the enriched GO-BP sets for the compared species and *N, M* their cardinalities. Each sum function in the numerator refers to one of the two GO-BP sets and aggregates the maximum similarities of its terms with the other set of terms. The aggregation of best matches between these two lists is averaged by dividing it with the sum of their sizes. The final distance matrix was defined by subtracting similarity scores by one (**Fig. 1B, step II**), and the phylogenetic clustergram was generated based on Ward’s minimum variance method (Ward, 1963).

### Phylogenetic analysis

Gene sequences of 16S and 18S rRNAs were used to construct the reference phylogenetic tree, as they traditionally portray the evolutionary proximities of species. The ClustalW tool (Sievers & Higgins, 2018) was used to quantify the pairwise distances and construct the final distance matrix, based on an *ad hoc* multiple sequence alignment (MSA). Furthermore, the amino acid sequences of HSP40 and HSP70 families were analyzed to examine their potential as surrogate, evolutionary markers. Consensus sequences for the HSP proteins were established, as heat shock proteins of the same molecular weight could vary significantly, even in the same organism. Each protein class consists of different members, which encompass certain, identical, functional domains, yet other additional components or their tertiary structure might be different. For instance, the human genome encodes 13 proteins of the HSP70 family and around 50 members of HSP40, which are divided in three main sub-families (Kampinga et al., 2009). Members of the same HSP family clustered to a consensus sequence pattern for each organism. Starting from the whole set of amino acid sequences, fragments were filtered out. The trimmed part fed the CD-HIT clustering algorithm with similarity threshold to 95% (Li & Godzik, 2006). CD-HIT keeps the longest sequence, as a representative feature of each cluster, conserving as much information as possible for each one. If the output included more than one clusters, then an extra step was performed, by constructing their multiple sequence alignment (MSA) with ClustalW and the respective hidden Markov model (HMM) with the HMMER3 hmmbuild algorithm (Finn et al., 2011). The final consensus sequence was generated using the hmmemit function of HMMER3. ClustalW was used to calculate the distance matrices of consensus sequences, similar to the case of ribosomal sequences.

To compare the phylogenetic dendrograms and evaluate their discriminative power, as well as their efficiency to reproduce well-shaped taxonomic clusters, organisms were projected on a two-dimensional plot, based on their distances. Specifically, the distance matrix of each phylogenetic approach was transformed into a two-dimensional orthogonal space, using the Multidimensional Scaling (MDS) technique (Borg & Groenen, 2005). MDS performs non-linear dimensionality reduction, projecting the data on a new orthogonal space, where distances among samples converge to the initial values, under a relative tolerance of cost function. The generated scatter plots illustrate the adjacency of species groups, indicating the divergence of each criterion through evolution.

### Identification of proteostasis components

To identify which biological processes, (as they are described on the standardized GO-BP graph), are significantly associated with proteostasis and how their profile changes across the three taxonomic kingdoms, we developed and implemented a two-steps clustering workflow (**Fig. S1.1**). The enriched GO-BP terms were first substituted by the respective sets of ancestral terms in the standardized graph. Then, the following workflow was applied to the three separate taxonomic kingdoms: i) Terms which were enriched in more than 90% of species were identified and filtered to reduce the semantic redundancy. Terms, whose descendants were also included in this set, were filtered out too. Hence, only uniquely defined terms remained in the list, namely the most semantically specific terms that were omnipresent in the examined PN semantic profiles. ii) Terms enriched in less than 90% of species were screened towards their ancestral paths to reveal their generic ancestors. As generic terms were designated all those having two edges distance from the graph root. If a generic term was identified to be linked with enriched terms in more than the 50% of kingdom species, then it was considered as significant. While these terms were not included in the initial enriched sets, the accumulation of enriched terms in their semantic branches made them indirectly associated with the PN semantic profile of many species in the same kingdom. The union of the term sets, derived from these two clustering steps, constituted the definite set of PN-related GO-BP terms, to represent the scaffold for the comparative analysis.

All generic biological processes, included directly or indirectly in the results of pathway analysis, for the majority of species entities in a taxonomic kingdom, were assumed as PN-related terms. Next, we quantified the association of species with these terms, using the results of the pathway analysis. For each PN-related term, the negative logarithm of its corresponding adjusted p-value or that of the minimum adjusted p-value of its descendant terms was kept, based on its inclusion or not in the results of pathway analysis, representing a reliable index of the enrichment of the given gene set in the PN of the examined organism. Finally, we decomposed the constructed association matrix, using the method of non-negative matrix factorization (NMF), in order to cluster the PN-related terms into three functional eigen-components and reveal the common and differential components of proteostasis across the different taxonomies.

### Evaluation of the classification performances of other evolutionary conserved mechanisms

GO-BP annotation was also exploited to inspect, similarly to PN, the clustering performance of 20 other fundamental biological processes, conserved across all analyzed species. Gene sets were retrieved from GO-BP for each species and biological process. Using these gene sets in pathway analysis could lead to significantly biased results (very high enrichment scores and artificially small p-values for the selected GO term, as well as its ancestral path). In order to mitigate potential bias infiltration due to the annotated content and the size of a specific gene set we introduced a randomization process to select the appropriate subset of genes of each pair of species and GO term (**Fig. S1.2**).

Specifically, the depth of genomic annotation and the distribution of hypergeometric p-values of the respective PN-related pathway analysis, were used as criteria to assert the optimal gene set size. If the annotation comprised less than 10 genes, then all of them were used as input for the pathway analysis, as this size is considered fairly small to generate critically biased results. For larger genomic sets, a random sampling procedure was implemented to generate different lists of genes, which sizes were selected, so as to produce approximately similar, extreme hypergeometric p-values, to those observed during the analysis of the respective PN-related gene set. An iterative binary search was employed to estimate the maximum subset of GO term annotation necessary for the pathway analysis, to yield the lowest log-transformed hypergeometric p-value to the order of magnitude of the average of the 10 lowest log-transformed p-values of the respective PN-analysis. As this procedure is based on the random selection of gene sets, the final solution is erratic, namely the optimal size changes based on the number and annotation of the selected genes. Thus, it was iterated 30 times, to create a distribution of optimal gene set sizes. Finally, different gene sets were generated with size randomly selected from this distribution, or equal to 10, in case the distribution mean was less than 10. The adoption of all these criteria by the workflow, attenuated the creation of either completely biased or non-informative semantic networks.

Prioritized pathway analysis of these sets of genes was implemented to obtain a semantic network profile for each species. If random gene sets were generated for a species based on the above workflow, then only the GO-BP terms enriched in more than 20% of the outputs were considered as part of the final semantic profile. Comparative analysis was used to calculate the final semantic distance matrix and its phylogenetic clustergram. Each phylogenetic tree was divided in three clusters, aiming to evaluate whether the examined biological process could reproduce the three taxonomic domains. Their efficiency was estimated with homogeneity and silhouette scores.

### PN contribution to other evolutionary conserved mechanisms

To derive semantic network profiles limited to the non-PN components of these biological processes and therefore assess the contribution of PN for taxonomic classification, PN-related terms were excluded from the semantic profiles of the examined conserved mechanisms. The comparative analysis was performed on these shortened profiles.

## Results

### Process implemented to evaluate evolution of the proteostasis network

As proteostasis maintenance is essential to sustain proper proteome functionality and cellular fitness, it represents *de facto* a conserved actor during evolution. We sought to study how the PN changes from prokaryotes to eukaryotes and to delineate the subsequent hierarchical tree, taking into account functional differences and commonalities recorded in its topology. As such, a common vocabulary was necessary to adequately describe the topology of the PN for all the species investigated. Thus, we used the Gene Ontology Biological Process (GO-BP) annotation to build the PN semantic profile in 405 species, based on gene lists associated with proteostasis. We also used the available rRNA nucleotide sequences and heat shock protein (HSP) amino acid sequences, to perform phylogenetic analyses on the same group of organisms, allowing therefore their comparison with PN-based trees. An exhaustive analysis of biological data public repositories (Ensembl (Cunningham et al., 2018), UniprotKB (The UniProt Consortium, 2018), and ENA (Leinonen et al., 2010)) was performed to collect the raw data. Organisms were included in the analysis by meeting the following criteria: i) genomic annotation in the GO-BP corpus; ii) availability of gene sequences of 16S (for Prokaryotes) and 18S (for Eukaryotes) rRNAs and iii) at least one annotated amino acid sequence of HSP40 (*dnaJ*) and HSP70 (*dnaK*). Using both manual and automated procedures (see 2. Materials & Methods), we generated comprehensive lists of rRNA, HSP40, HSP70 sequences and a gene list related to proteostasis for 405 organisms (93 Eukaryotes, 250 Bacteria and 62 Archaea; **Fig. 1A, step II**). Pathway analysis was used to translate each gene list into GO-BP terms, which imprinted the PN-associated networks (**Fig. 1B, step I,** for gene lists and pathway analysis results see **Code and Data Availability** section). A phylogenetic comparison of these functional profiles was performed through the calculation of their semantic similarities. Specifically, we used a standardized version of GO-BP in conjunction with semantic operators, to quantify the similarities of term lists and create a phylogenetic dendrogram by clustering together species with high semantic similarities (**Fig. 1B, step II**). Such clustergram entailed the use of a reference version of GO-BP, for the calculation of the inter-species topological similarities, neutralizing annotation imbalances which incurred bias in the description of proteostasis among different organisms. This was achieved by the exclusion of terms which reside low in densely populated ontological branches, through the application of cutoff criteria for two metrics, Information Content (IC) and Semantic Value (SV). Following this method, a pruned graph of 2693 terms was generated (**Fig. 1C**). Finally, the PN-based hierarchical tree, as well as those of rRNA and HSP sequences, were built using agglomerative clustering (Ward, 1963).

### Ribosomal RNA, HSP40, HSP70 and PN-based phylogenies

Clustergrams (**Fig. S2.1-4**) were constructed to compare rRNA nucleotide sequences, HSP40, HSP70 amino-acid sequences and the semantic topology of PN profiles. To further explore the characteristics of each evolutionary tree, species were projected on a two-dimensional space (**Fig. 2A**) using the Multidimensional Scaling (MDS) algorithm (Borg & Groenen, 2005). Both rRNA sequences and PN semantic profiles produced nearly independent sub-groups for each established taxonomic domain. The only inconsistency of rRNA-based phylogenetic clustergram with the reference was observed with a group of *Streptophyta*, which clustered close to Bacterial species (**Fig. S2.1**). In a related way, regarding the PN-based classification, seven Bacteria (mainly *Planctomycetes*) were embedded into the branch of archaea (**Fig. S2.2**). PN evolution appeared more constrained than that of rRNA, which led to distantly separated kingdoms. The distributions of PN- and rRNA-based distances corroborated this finding (**Fig. S2.5-6)**, as the PN semantic profiles produced systematically lower inter- and intra-distances among the taxonomic domains. This probably reflects the heterogeneity of the PN content, bisected into taxonomy-specific components and others that are conserved across evolution. Nevertheless, the pairwise distances among species for rRNA and PN showed high correlation (**Fig. 2B**). Concerning the HSP-derived classification, a poor correlation of the HSP40 and HSP70 sequences with evolution was observed, as they only succeeded to separate eukaryotic and prokaryotic kingdoms, even so not flawlessly. A weaker but not negligible correlation with rRNA sequence-based distances suggests that the intra-distances of taxonomic clusters follow approximately the same distribution to their inter-distances (**Fig. S2.7-8**). These observations imply that HSPs, which are individual components of the PN, lack informative power as a marker of species evolution.

**Figure 2:**
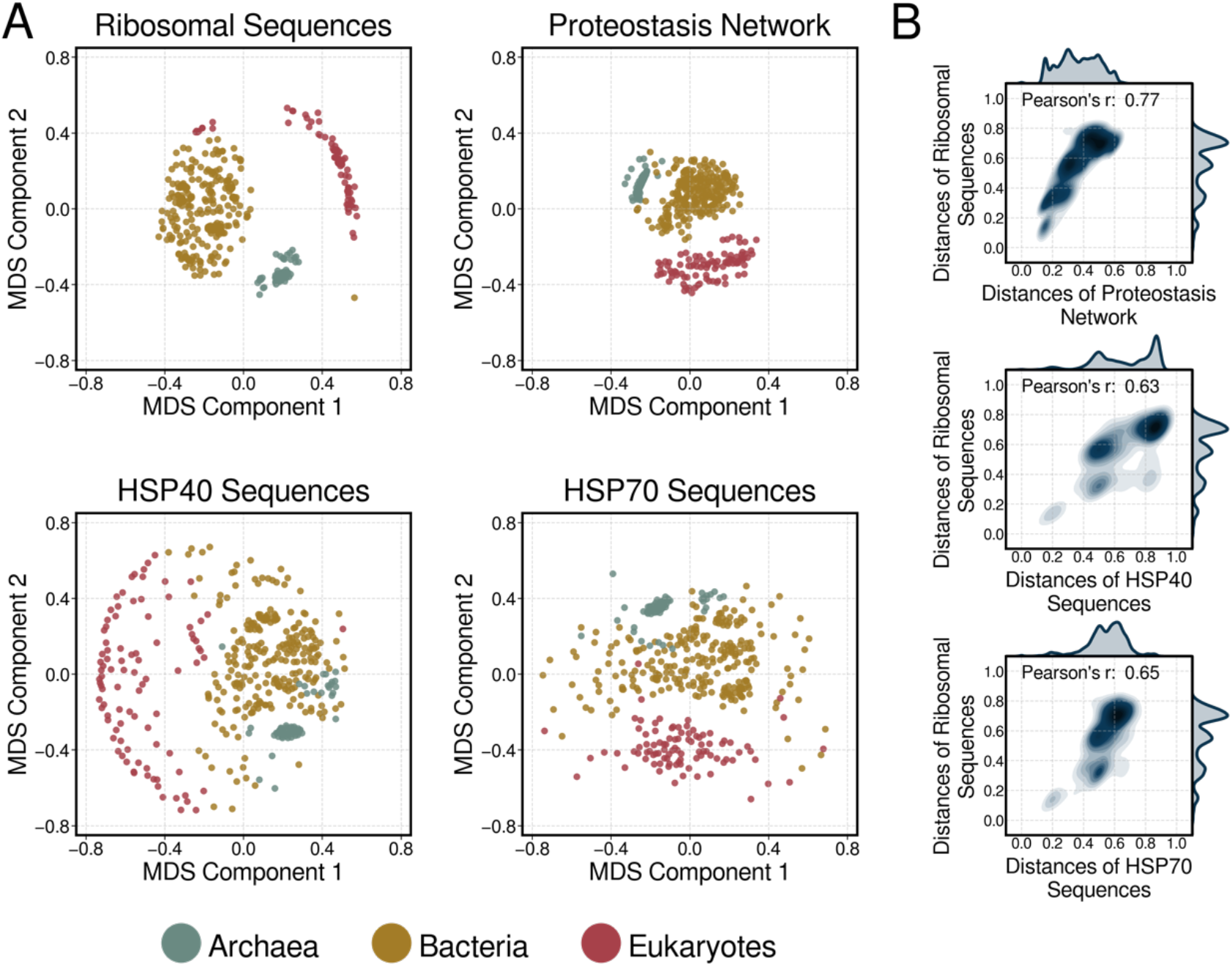
Establishment of PN-based phylogenetic trees. **(A)** Two-dimensional representation of organisms in function with their taxonomy. The derived evolutionary distance matrix of each criterion (rRNA, proteostasis and HSPs), was transformed into a 2-dimensional orthogonal space through the Multidimensional Scaling (MDS) algorithm, reflecting the two larger, dimensions of observed variation (termed eigen-components). Similarity distances of each organism from the centroid of the single class problem are projected in those exploratory scatter plots. **(B)** Pearson correlation of pairwise distances of rRNA sequences with the other three measures. Correlation is unbiased from taxonomic domain sizes, as we used 80 randomly selected species from each domain for the calculation.

We next sought to compare the accuracy of PN, as well as rRNA and HSPs sequences to classify species of the same taxonomic domain. As such, we examined the Class-level categorization of Archaea and that of Phylum-level for Eukarya and Bacteria. For each criterion, different clustering models of species, for a range of predefined number of clusters, were generated and the consistency of each model with the reference classification was assessed, using the homogeneity score (see Appendix). The output corroborated the findings inferred from the rRNA sequences and revealed an overall homogeneity of PN profiles in Bacteria and Archaea (**Fig S2.9**). None of the criteria succeeded to impeccably reproduce the number of clusters of the reference taxonomic groups, indicating that species of different lower-level taxonomies share similar profiles. In general, the results confirm the appropriateness of the rRNA sequence homology as an insightful, evolutionary measure. Regarding the PN-based clustering models, the low homogeneity scores for the prokaryotic domains demonstrate high conservation of PN semantic profile in each domain, regardless of the lower-level taxonomic classifications. In contrast, the accuracy of PN-based models was significantly improved for the case of Eukaryotes, suggesting that a key avenue of their evolutionary adaptation resides in the diversification of the plasticity of their genetic circuitry, enabling novel, emergent cellular functions. Hence, PN encompasses variations that coherently segregate eukaryotic species, rescuing phyla segregations to a large extent. HSP-derived clusters were similar to those obtained with rRNA sequences in Prokaryotes, but declined among Eukaryotes, probably due to the high variation of protein families in species-specific profiles. To sum up, the efficiency of PN as a taxonomic marker in the lower evolutionary taxonomies is inverse to that of HSP-sequences, implying that the profile of proteostasis is more informative for complex organisms, while lower species have significantly similar PN semantic topologies.

### Proteostasis as a functional metric to trace evolution

The detection of quasi-omnipresent GO-BP terms in the PN semantic profiles and their agglomerative clustering, led to the identification of 50 PN-related semantic groups (generic biological processes) and the quantification of their association with the examined species (**Fig. 3A**). The NMF-based transformation of the association matrix into a matrix of three eigen-components (**Fig. S2.10**), each showing a noticeable kingdom-specificity (**Fig. S2.11**), revealed the conserved, as well the differential parts of PN-profiles across the three kingdoms. Metabolic processes (both catabolic and biosynthetic), transport- and localization-related processes, protein folding and cellular responses, especially due to temperature stimulus, were enriched in all tested species, thereby representing the PN “conserved core”.

**Figure 3:**
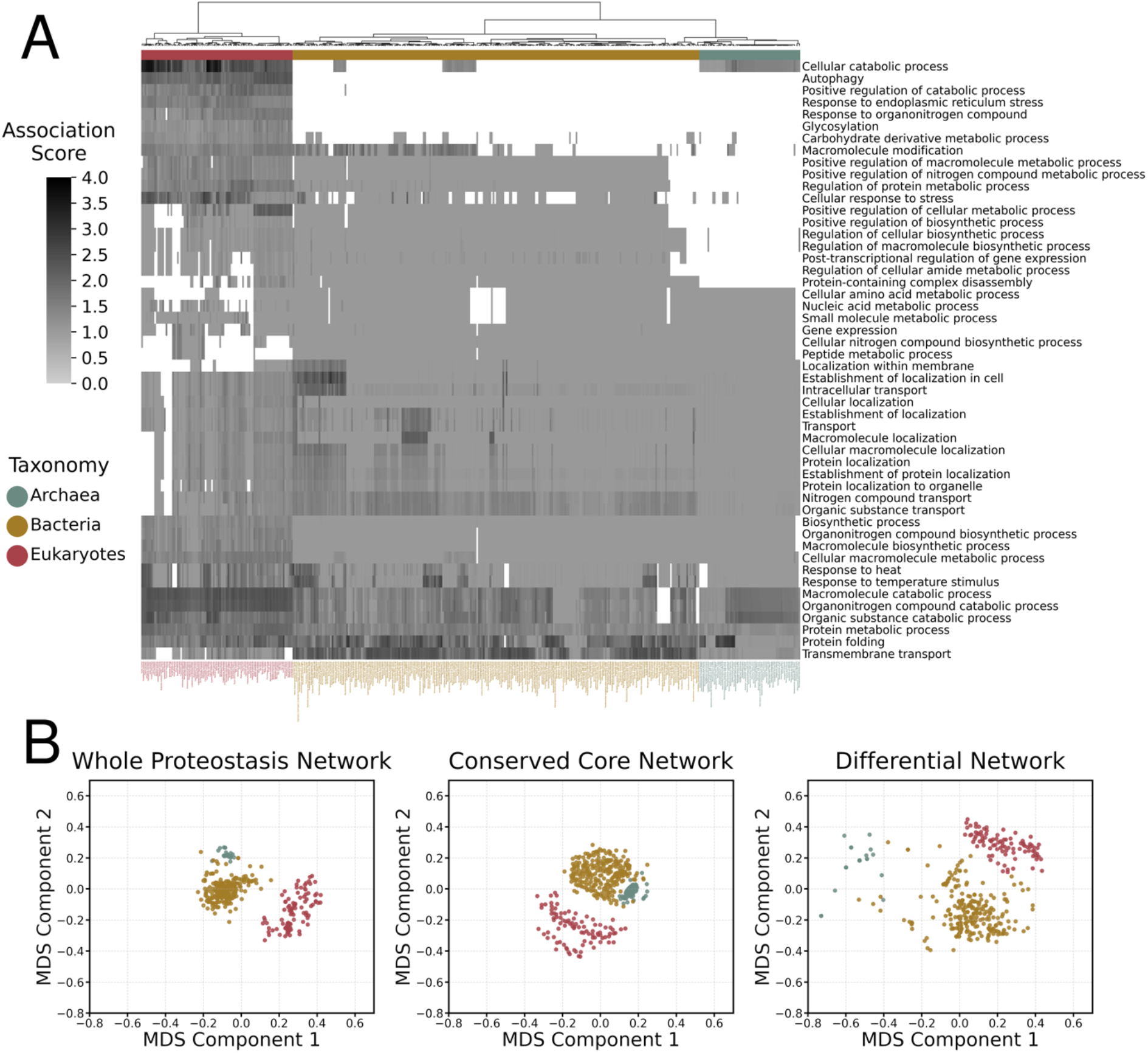
PN composition evolution throughout various phyla. **(A)** The association matrix of proteostasis-related semantic groups with the examined species. The score of each semantic group corresponds to the average negative log-transformed adjusted p-value of the GO-BP terms, which have been clustered into this group. **(C)** Two-dimensional representation of species based on their proteostasis semantic profiles. Conserved and differential components were used separately to investigate their contribution to taxonomic domain separation.

The obtained PN-related term groups presented significant semantic relevance or even overlap. For example, the “conserved core” included many terms which correspond to catabolic processes. All these generic terms have been identified because different components of catabolism were found enriched for all the PN-related gene sets. Protein catabolism is one of these components. However, the identification of “protein metabolic process” in the “conserved core”, signifies that additional processes related to the modification of protein in cell have been found in the PN semantic profiles (such as “protein maturation”, “proteolysis”, “formation of translation preinitiation complex”, “protein peptidyl-prolyl isomerization” and “histidyl-tRNA aminoacylation”), apart from “protein catabolic process”. Two other biological processes, highly represented in the “conserved core”, are localization and transport. Additionally, terms related to response to stress designate another distinct set of processes. “Response to temperature stimulus” is the semantic parent of “response to heat”, so their co-existence in the “conserved core” produces semantic redundancy. This result has been caused due to genomic annotation resolution imbalances, across species of different taxonomies. Particularly, “response to heat” has been found as a quasi-omnipresent term in Archaea, while “response to temperature stimulus” has been detected in the PN profiles of Bacteria and Eukaryotes. Both of them imply the association of proteostasis with components of cellular response to abiotic stimulus.

The “conserved core” includes the main PN-profile of Archaea (eigen-component 3). This profile lacks regulatory mechanisms, which have been assigned to that of Bacteria (eigen-component 1) and Eukaryotes (eigen-component 2). While prior knowledge about the regulatory processes in Archaea exists and has been integrated within the genomic annotation of GO-BP, this finding implies that they do not have a strong contribution to proteostasis machinery. For instance, the concept of “post-transcriptional regulation of gene expression” exists in both Archaea and Bacteria. Its genomic annotation in Archaea contains translation initiation (*aif5A, tef5A, tif5A, eif2g, eif5a*) and elongation (*efp*) factors, synthases (*dph2, dph5, dph6, dphB*), reductases (*cbiJ*) and ribosomal proteins (*rpl1, rpl13, rpl1P, rpl1p, rplA, rps4*), which participate in the process of translation and protein biosynthesis. On the other hand, the same regulatory process is neatly equipped in Bacteria, containing genes, which regulate the translation and consequently could be considered as molecular entities of proteostasis apparatus, such as small heat shock proteins (*ibpA*), ribosome hibernation factors (*hpf*), elongation factors that act under stress conditions (*lepA*) and proteins which monitor the translation of Sec system components (*secM).* Hence, the expression of such regulatory entities forms a more complex PN semantic profile for Bacteria compared to that of Archaea.

In addition, specific processes related to programmed cell death and signaling pathways were enriched in Eukaryotes. Enrichment in terms associated with the response to endoplasmic reticulum (ER) homeostasis imbalance was effective in all examined eukaryotic species, as expected, pointing out this compartment as a hotspot for proteostasis. Regarding the process of autophagy, many descendant terms identified exclusively in Eukaryotes, related to either associated membranous structures (“autophagosome assembly”) or subprocesses and mechanisms (“lysosomal microautophagy”, “autophagy of nucleus”, “chaperone-mediated autophagy”). The extended PN profiles of Eukaryotes were further enriched by the integration of protein glycosylation and additional regulatory and cellular response mechanisms.

We next used the association of species with each GO-BP group, to derive their two-dimensional representations for the whole PN (**Fig. 3B**, left), the common PN (**Fig. 3B**, middle) and the differential PN (**Fig. 3B**, right). PN profiling produced independent sub-groups for each taxonomic domain, as it was expected from the previous results. These sub-groups increased their density when analyzing the PN “conserved core” (**Fig. 3C**, middle) and seemed sparser when analyzing the differential PN (**Fig. 3C**, right). In the case of Archaea, the great sparsity in the differential PN reflects maximum semantic distances among species, which have been caused by the removal of the PN “conserved core” and the subsequent elimination of their semantic profiles. Collectively, these data unveil conservation and divergence within the PN, highlighting circuits of ubiquitous functions and emerging mechanisms indicative of each taxonomic domain.

### Impact of PN evolution on other functional networks

To evaluate the robustness of our methodology and to depict the association of the PN with other cellular processes, we examined the semantic profiles of other evolutionary conserved mechanisms. The ability of each mechanism to separate the main taxonomic domains was evaluated using the homogeneity (HS) and silhouette (SS) scores (see Appendix). To this end, we analyzed 20 conserved processes, as defined in the GO-BP, and both semantic and phylogenetic analysis were performed based on their genomic annotation. Then, GO-BP terms included in those semantic profiles, which were also related to the obtained PN semantic groups, were excluded to repeat the phylogenetic analysis upon their exclusion. This showed that biological processes, which emerge as gains of functions during species evolution, or those, which evolved through higher functional complexity, segregate sufficiently the three kingdoms (**Fig. 4; S2.12A-S2.31A**). The biological processes identified through this approach are related to cell compartmentalization, regulatory networks, lipid metabolism and DNA recombination. Some performed marginally better in terms of accuracy, as taxonomic metric, compared to proteostasis or to rRNA sequences - i.e. slightly higher HS values due to the erroneously classified Bacteria in the case of PN, and the group of plants in the case of rRNA sequences. Nevertheless, the phylogenies of these biological processes exhibited lower SS values than those obtained for PN (**Fig. 4B**), which means that the produced clusters were sparser (especially within Prokaryotes) and consequently, the phylogenetic trees contained broader clades. In addition, parts of the aforementioned processes or those with narrower functional networks showed weaker performance, concerning clustering efficiency, especially due to their strong commonalities among the Prokaryotes. HS measurements were below 0.8, because many Archaeal species were classified in Bacteria, and *vice versa.* To evaluate the contribution of the PN to those machineries, we artificially removed proteostasis-related components from each semantic profile and measured the impact of such action on HS and SS (**Fig. 4; S2.12B-S2.31B**). Only a few processes (“cellular component assembly” and “lipid metabolism”) retained adequate information to distinguish accurately the taxonomic domains, implying that their mechanistic frameworks can differentiate the taxonomic kingdoms. In general, all the processes with accurate performance suffered from low silhouette scores, and some lost their phylogenetic congruity. This analysis indicates that a significant part of the semantic description and components of PN is included in other mechanisms involved in cell homeostasis and functions.

**Figure 4:**
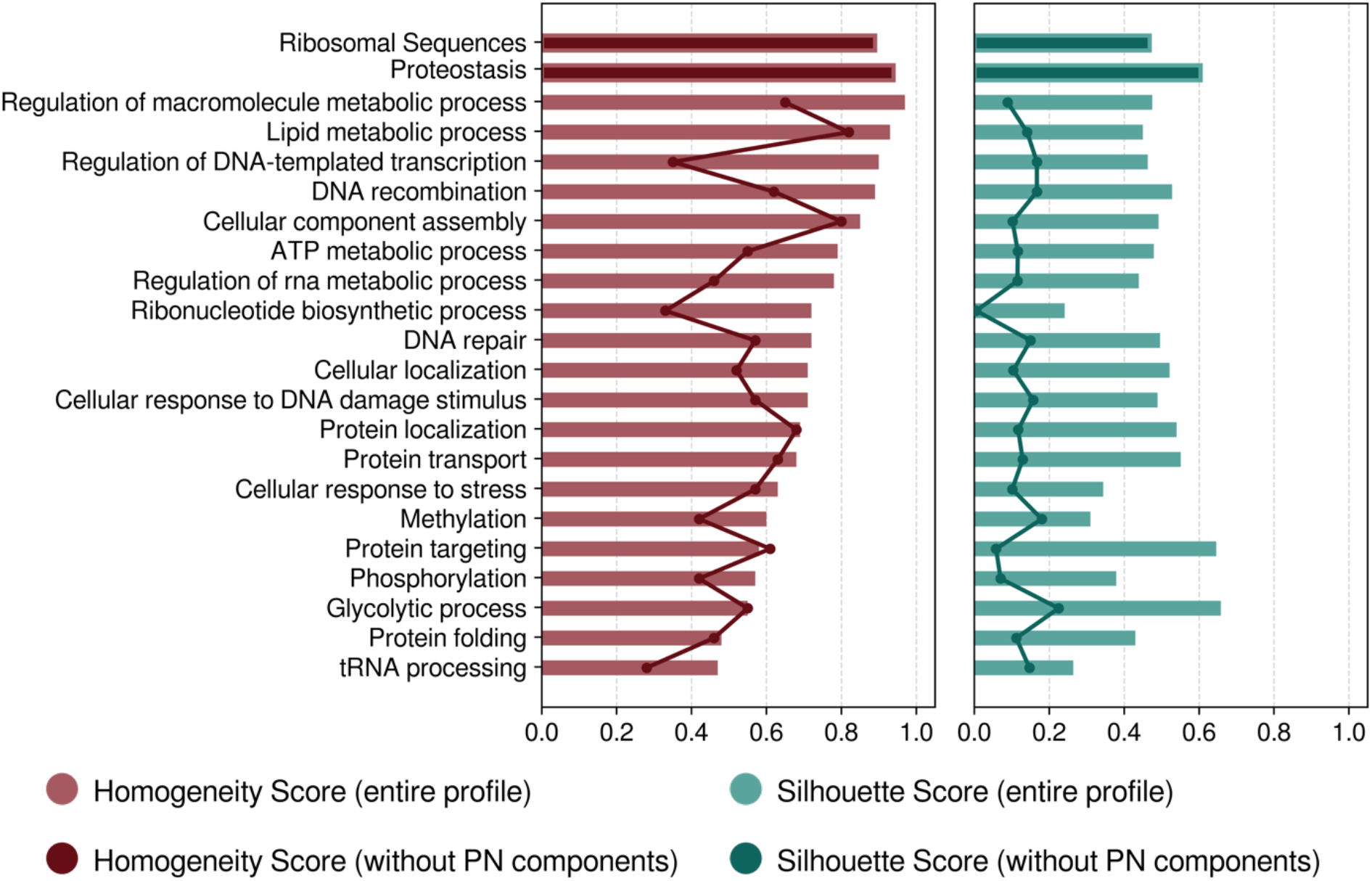
PN contribution to other evolutionary conserved mechanisms. Homogeneity score (**A, red**) refers to the separating ability of each biological process regarding the three main taxonomic domains, through the respective semantic network, derived from the pathway analysis of related genes. Silhouette score (**B, green**) indicates the degree of cohesion of cluster inference, by measuring the trade-off between intra- and inter-distances of each cluster member. Bars display these scores for the entire machinery of each process whereas the solid lines illustrate the same scores calculated after the removal of proteostasis related components from the semantic profile of each process.

## Discussion

Taxonomies based on molecular sequences have increased our understanding of evolutionary processes. Phylogenies based on isolated macromolecules conserved among evolution, such as nucleic (DNA/RNA) or protein sequences, have their limits as they do not accurately represent the complexity of life evolution. In this study, we designed and applied a novel approach for phylogenetic comparison which uses the semantic graph as a new metric, to evaluate the complexity of protein homeostasis through monitoring the proteostasis network (PN). To the best of our knowledge, it is the first time that semantic network analysis is used to define an evolutionary marker. Regardless of the ability of the PN topology to indicate the main taxonomic characteristics of a species, the proposed approach stands as a novel strategy for taxonomic classification. It relies on the semantic comparison of ontological graphs, and the quantification of their divergence across different species, rather than the analysis of individual sequences. Using a standardized ontological framework, the semantic interpretation of gene/protein sets for different species provided biological insights about the impacts of the evolutionary pressure, and the extent of conservation of mechanisms among Eukaryotes, Archaea and Bacteria. In this premise, our study aimed at illustrating the complexity of the modular architecture of proteostasis. In addition, our analysis measured species-specific topological differences, and translated them into evolutionary branches. We show that monitoring PN characteristics provides reliable information about the evolution of living organisms in the level of kingdoms, whereas monitoring its individual components has limited interpretation.

First, PN data succeeded to separate the three main taxonomic domains almost infallibly, performing as accurately as rRNA sequences do. Moreover, PN performed much better than isolated PN components (conserved families of heat shock proteins, **Fig. 2**). This primarily indicates that the proposed new method is able to disclose evolutionary differences among species of different taxa. It also demonstrates the utility of PN as a reliable evolutionary marker, able to classify species according to their main taxonomy, contrary to the limitations of the sequence-based approaches. The efficiency of the PN metric to increase taxonomic resolution dropped in Bacteria and Archaea, likely due to the deficiency of the proposed method to adequately quantify nuances in the respective semantic topologies. GO-BP provides a descriptive genomic annotation which might not be suitable to elucidate slight functional differentiations between species. However, the impediment is that at the moment, there is not any controlled vocabulary, which describes the universe of cellular functionality at the level of pathways’ topology for thousand species. For instance, the BioCyc database (Karp et al., 2019) contains curated and computationally inferred annotation for thousands of species only for metabolic pathways, while Reactome Pathways (Jassal et al., 2020) provides curated annotation for the network of pathways of a few model-species. A way to resolve this issue could be the massive integration of data from different vocabularies into a unified, ontological schema. In addition, the weaker segregating capacity of the PN in the two prokaryotic kingdoms might also imply that their evolutionary pressure was managed through the functional diversification of selected genes (e.g. mutagenesis) or chromosome variations. In Eukaryotes, the complexity and adaptability of molecular circuitries became a major driving force. For the Eukaryotes, the evolution of compartmentalization impinges on the complexity of protein circuitry, prompting evolution of the PN to cope with these additional constraints (Powers & Balch, 2013).

Deconvolution of the PN in various molecular sets revealed that many, if not all cellular processes, are connected to the PN, especially for Eukaryotes. The obtained semantic profile of Eukaryotes contains the majority of systemic processes that delineate the proteostasis network of *Homo sapiens* (The Proteostasis Consortium, Overall coordination et al., 2022). The conserved PN component across the vast majority of species is linked with protein production, folding and degradation. It is also linked with responses to external or internal stimuli, activation or repression of anabolic and catabolic processes, to maintain cell homeostasis, as well as the proper localization of macromolecules. This comprises heat shock proteins, which perform poorly in terms of separating the taxonomic kingdoms, but were recently shown to organize beyond the “de novo versus stress-inducible” scheme, into a layered core-variable architecture in multi-cellular organisms (Shemesh et al., 2021). In addition to this, in order to cope with proteome complexity that arises with evolution, conserved core chaperones increased in abundance and new co-chaperone families appeared (Rebeaud et al., 2021). Other functional modules, considered as ‘gain- or loss-’ of adaptive functions, were also found to be connected to the PN. This, in turn, could help identify the role of PN in maintaining those functional changes (**Fig. 4**). The instrumental role of proteostasis as a robust indicator of cellular and organismal adaptation to evolutionary cues, is highlighted by the taxonomic underperformance that the other mechanisms linked to PN exhibit when PN components are excluded (**Fig. 4**). This provides evidence for a tight coupling between proteostasis and other major biological processes. As such, the architecture of the PN encapsulates critical biological information to categorize species according to their complexity and acts as a clear-cut fingerprint of evolution, as it has co-evolved with the cell proteome and provided a driving force for adaptation to favor emergence of new traits.

Encouraged by the finding that PN sets a novel and reliable evolution metric for the main taxonomies, we feel tempted to propose the investigation of the use of this integrated information, as a quantitative trait to categorize diseases, and possibly their treatments. Approaches relying on the analysis of PN sub-networks (e.g. the “chaperome”) were proposed to be effective in various pathologies, such as cancers or degenerative diseases (Brehme et al., 2014; Hadizadeh Esfahani et al., 2018; Taldone et al., 2014), or in cell differentiation (Vonk et al., 2020). Considering that the “chaperome” represents an evolutionary conserved part of the PN, one might envision an approach relying on the PN, as defined in our study, to assess how its deregulation could mark disease appearance, evolution, or even propose specific nodes as potential therapeutic targets. At last, at a time of an unprecedented world pandemic with SARS-CoV2, one might also consider PN evolution, and its quantitative measurement, as a tool to predict the evolution of human and animal pathogens. The evolution of influenza virus is affected by the targeted alteration of the host’s cells PN, mostly through perturbations of HSF1 and HSP90 expression, once again belonging to the conserved core of the PN (Phillips et al., 2017). It may illustrate the needs for any pathogens to rely on a given host’s PN, which could be evaluated using molecular data sets (e.g. protein-protein interactions, gene expression). Our results could pave the way to investigate PN alterations and their consequences for the physiology and pathology of their hosts.

## Supporting information

Koutsandreas_2023_SUPPL

## Appendix

### Gain Ratio, Homogeneity and Silhouette Scores

Gain Ratio (Han et al., 2011) is used as a feature selection measure in data mining. Given a dataset *A*, with samples belonging to a set of classes *C* = {*c*_1_, *c*_2_,…, *c_N_*} and *D* a subset of *A*, the entropy of the classes’ distribution in subset *D* is defined as follows:

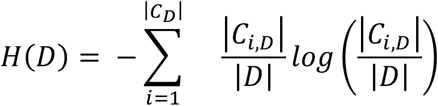

where |*C_i,D_*| is the number of samples in partition *D*, which belong to class *C_i_*. *H*(*D*) quantifies the expected information, to classify correctly a sample *a_i_* in *D*. This entropy is maximized when each sample belongs to a different class. During the training process of decision trees, discrete or continuous variables (features) are examined to separate samples, with respect to their classes. Assume that a feature *F* is selected to separate further the samples in *D*, producing *M* groups. Then, the entropy of *D* is re-calculated, taking into account that partitioning:

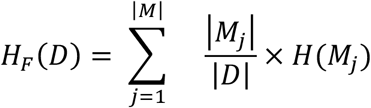

which is the weighted mean entropy of the derived *M* subsets. Information Gain of feature *F* is equal to the reduction of entropy after the split:

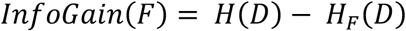

Another useful measure is the Split Entropy, which is defined as the derived uncertainty due to the partitioning of samples:

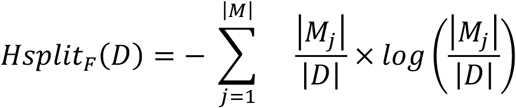

Split Entropy value increases in function with the amount of the produced subsets and it is maximized when feature *F* creates a novel branch for each sample. It could function as a penalty factor, in order to avoid the selection of features, which tend to segregate the dataset into numerous clusters. Gain Ratio uses that factor to normalize the Information Gain, providing an unbiased measure of splitting information:

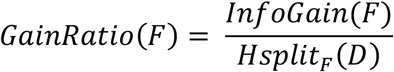

Homogeneity (HS) (Rosenberg & Hirschberg, 2007) and silhouette (SS) (Rousseeuw, 1987) scores evaluate specific properties of an unsupervised clustering outcome, given the true classes of the data samples and their pairwise distances. Assuming that the samples of dataset *A* are being classified into a set of clusters *K* = {*k*_1_, *k*_2_,…, *k_m_*} by an unsupervised clustering algorithm. The number of samples of class *i* which are assigned to cluster *j* is denoted as |*A_ij_*|. HS evaluates the quality of the derived *K* clusters to contain objects belonging to a unique class, by measuring the conditional entropy of the classes’ distribution given the proposed clustering:

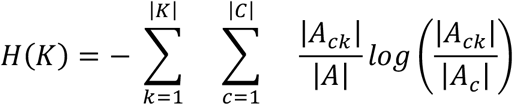

Formally, *H*(*K*) quantifies the uncertainty about the distribution of samples in the set of *C* classes, given the *K* clusters. If each cluster *k* is homogenous (i.e. it contains samples from only one class), then the conditional entropy *H*(*K*) is equal to zero. The conditional entropy is maximum (equal to *H*(*A*) - the entropy of the classes’ distribution in *A*) when the proposed *K*-clustering does not provide any information about the real classification. Homogeneity score is defined as:

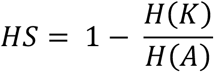

Using both the normalized entropy and the subtraction, HS is bounded in the range [0, 1] and its desirable value is equal to 1. Silhouette score is a measure of cluster cohesion membership, as it quantifies the trade-off between intra- and inter-distances of the derived clusters. Given the aforementioned *K*-clustering, for each sample *i* assigned to cluster *k_n_*, the factors *f*_1_(*i*) and *f*_2_(*i*) are defined as follow:

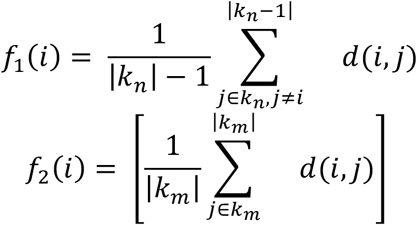

where *d*(*i,j*) is the distance between samples *i* and *j*. The factor *f*_1_(*i*) is the average distance of sample *i* to the other samples in cluster *k_n_* and indicates the merit of the assignment to that cluster. The factor *f*_2_(*i*) is the minimum average distance of sample *i* to all samples in any other cluster, apart from cluster *k_n_*. Namely *f*_2_(*i*) measures the inter-distance of sample *i* to its neighboring cluster. The silhouette coefficient of sample *i* is defined as:

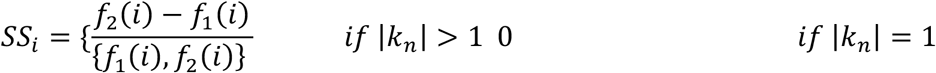

*SS_i_* ranges from −1 to +1, where a high value implies that *i* is well-located in its group and far from the other clusters (without considering if it is correctly classified). The mean silhouette over all data samples measures the average cohesion of the derived clusters:

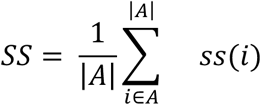

## Acknowledgements

This work was funded by grants from INCa (PLBio), FRM (DEQ20180339169) and ANR (ERAAT) to EC; from INSERM (International Research Project – TUPRIC), EU H2020 MSCA RISE-734749 (INSPIRED) to AC and EC; from ANR (ANR-15-CE12-0003-01) and FRM (DBF20160635724) to BF and ELIXIR-GR (MIS 5002780) to TK.

## Author Contributions

AC and TK developed and implemented the computational approach, BF and EC provided the biological conceptual framework to the analyses. All authors worked on the manuscript.

## Author Contributions

EC is a founder of Thabor Therapeutics. AC is founder and CEO of e-NIOS Applications PC.

## Code and Data Availability

The main part of code written in support of this manuscript is publicly available on a GitHub repository at https://github.com/thodk/proteostasis_imprinting_across_evolution. Some tasks were performed using the software of e-NIOS Applications PC, so the respective code is not available. However, an open version of BioInfoMiner and that of semantic comparative analysis are available in the Galaxy application http://www.biotranslator.gr:8080 upon registration. All the results and the data used to produce them, are available in a figshare collection at https://figshare.com/projects/Proteostasis_imprinting_across_evolution/113946.

## Notes

https://github.com/thodk/proteostasis_imprinting_across_evolution.

http://www.biotranslator.gr:8080

https://figshare.com/projects/Proteostasis_imprinting_across_evolution/113946

### Summary of Updates

Edits were made in the text for more clarity

