## Supplementary material for "Protein homeostasis imprinting across evolution": Koutsandreas_2023_SUPPL

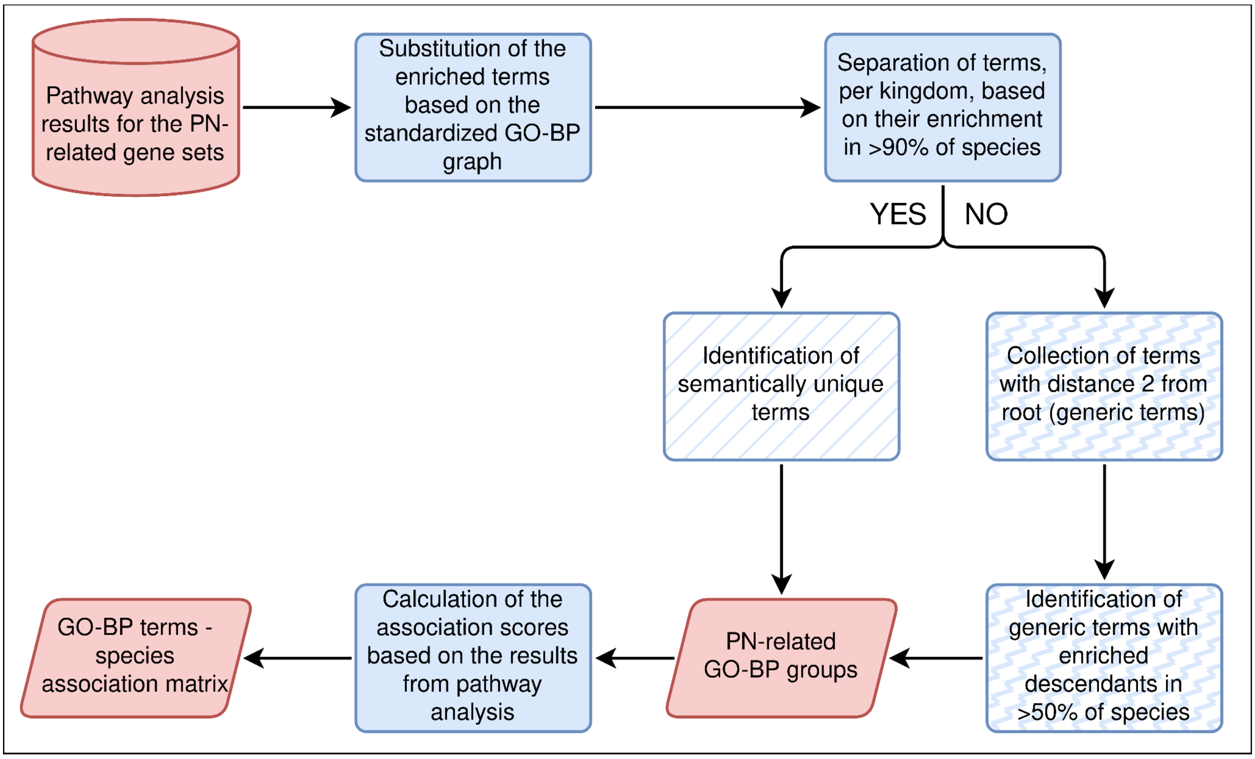


**Figure S1.1**: Schematic representation of the developed workflow for the identification of PN-related semantic groups, based on the results of pathway analysis.

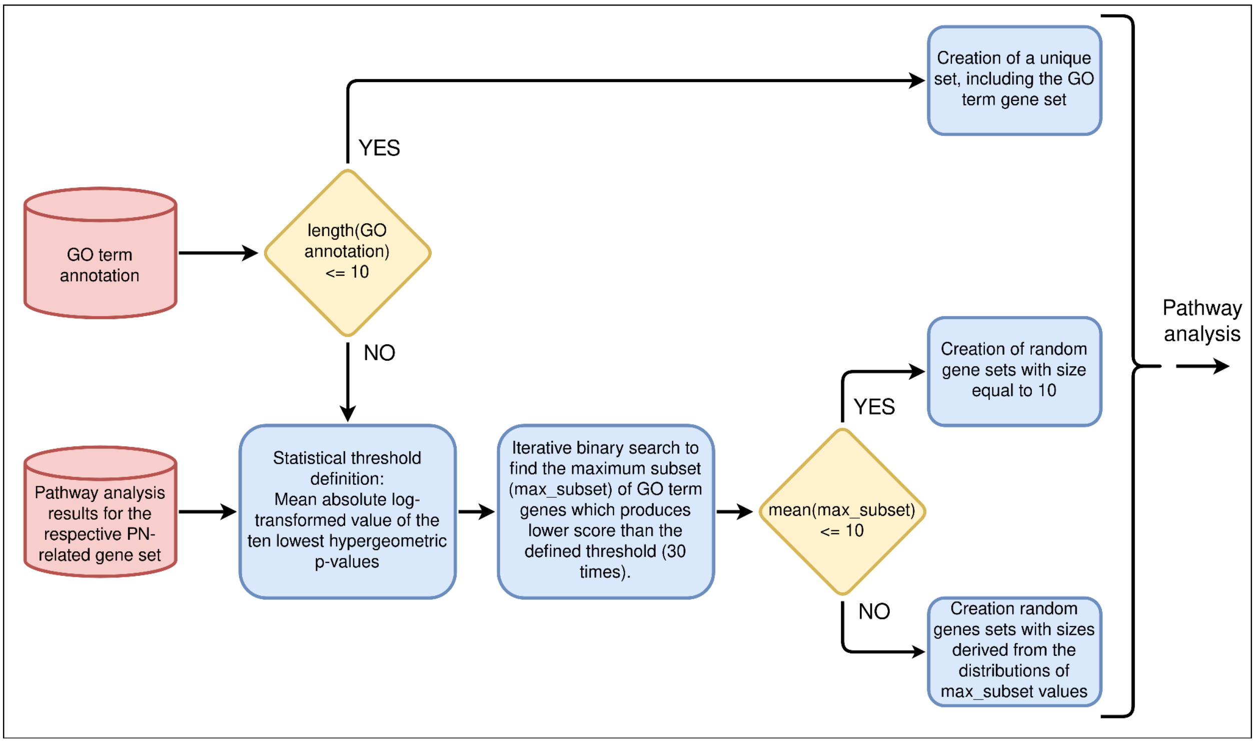


**Figure S1.2**: Schematic representation of the developed workflow for the determination of gene sets, used to derive the semantic profiles of the selected conserved mechanisms, through pathway analysis.

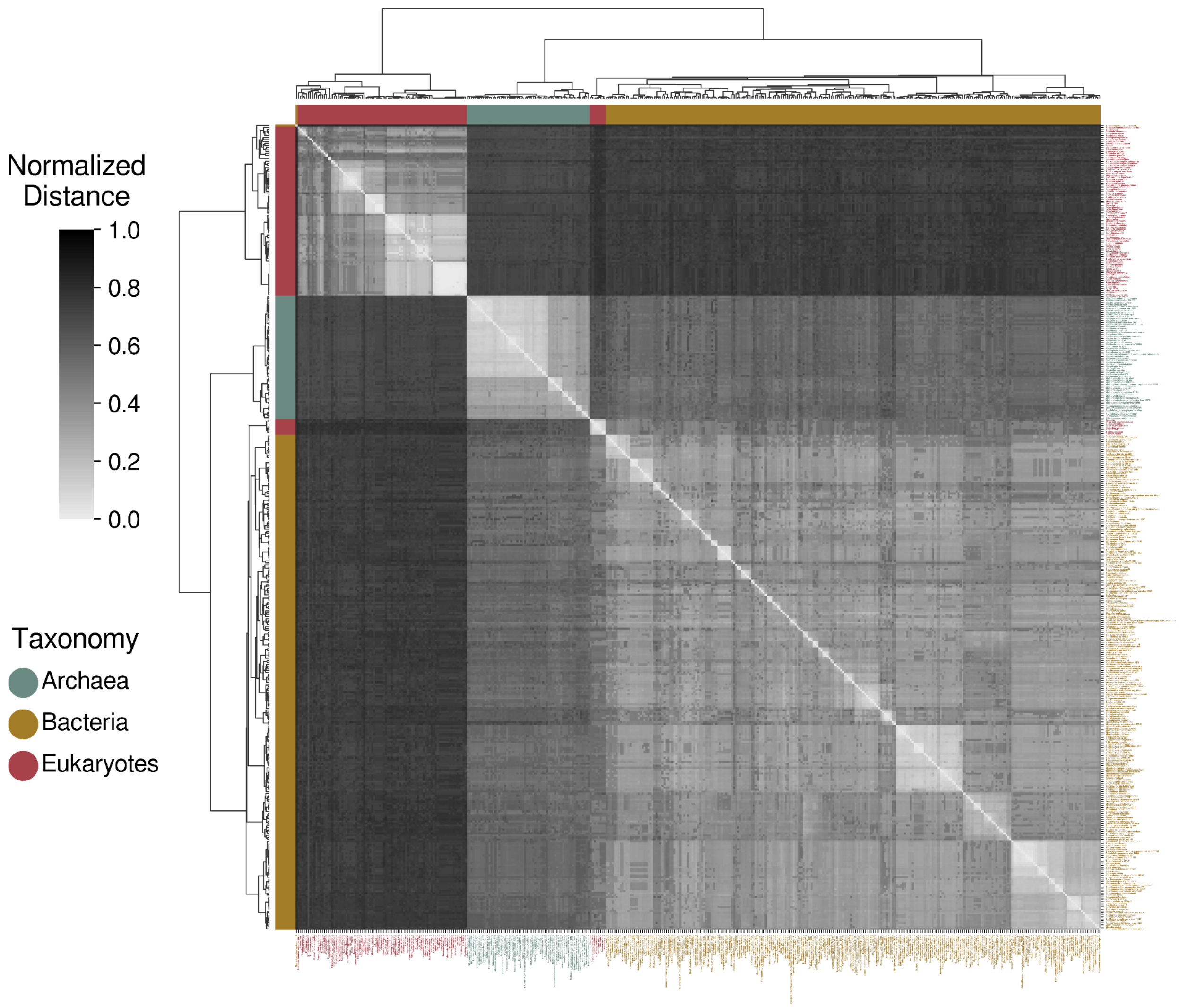


**Figure S2.1**: Phylogenetic clustergram derived from the comparison of the genomic ribosomal sequences (18S for eukaryotes and 16S for prokaryotes), using the ClustalW algorithm (Sievers & Higgins, 2018).

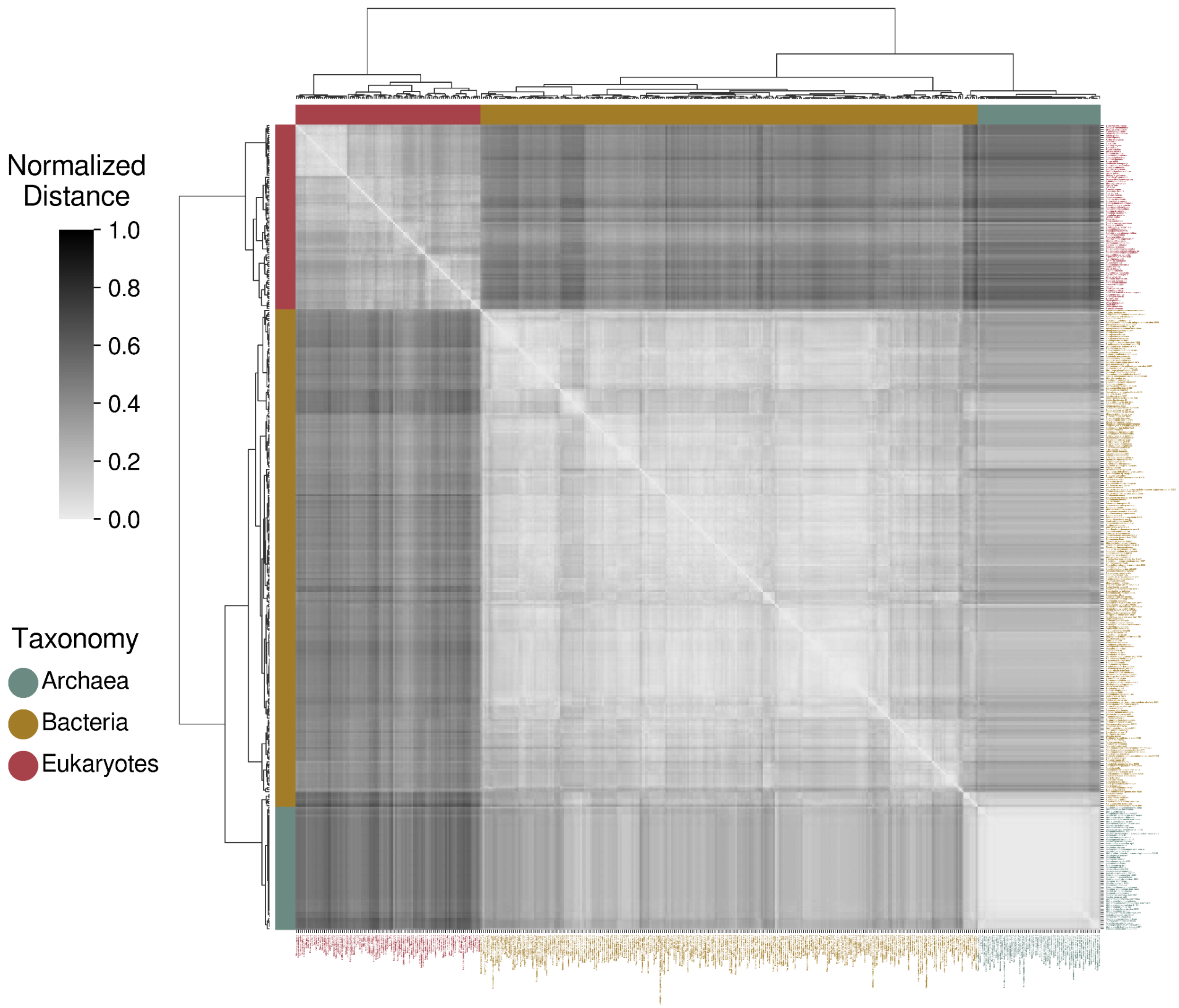


**Figure S2.2**: Phylogenetic clustergram derived from the comparison of proteostasis semantic networks. The networks were constructed performing pathway analysis on proteostasis related gene sets, using the GO-BP domain (The Gene Ontology Consortium, 2019). Semantic analysis operators were applied to compare the networks based on the GO-BP graph topology.

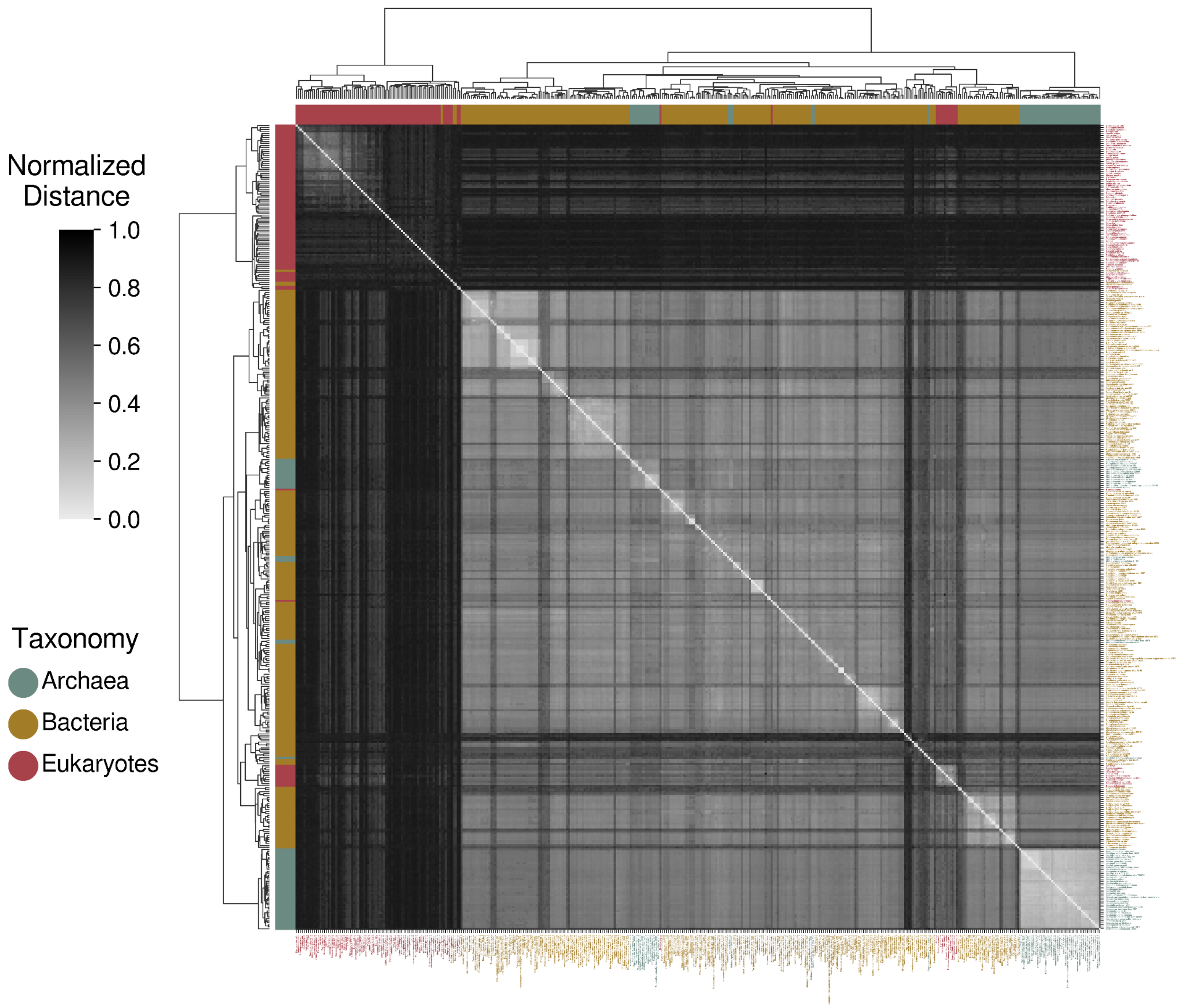


**Figure S2.3**: Phylogenetic clustergram derived from the comparison of heat shock protein 40kDa (HSP40) sequences. CD-HIT (Li & Godzik, 2006) and HMMER3 (Finn et al., 2011) were used to construct a consensus amino acid sequence for each species. The phylogenetic comparison was performed using the ClustalW tool (Sievers & Higgins, 2018).

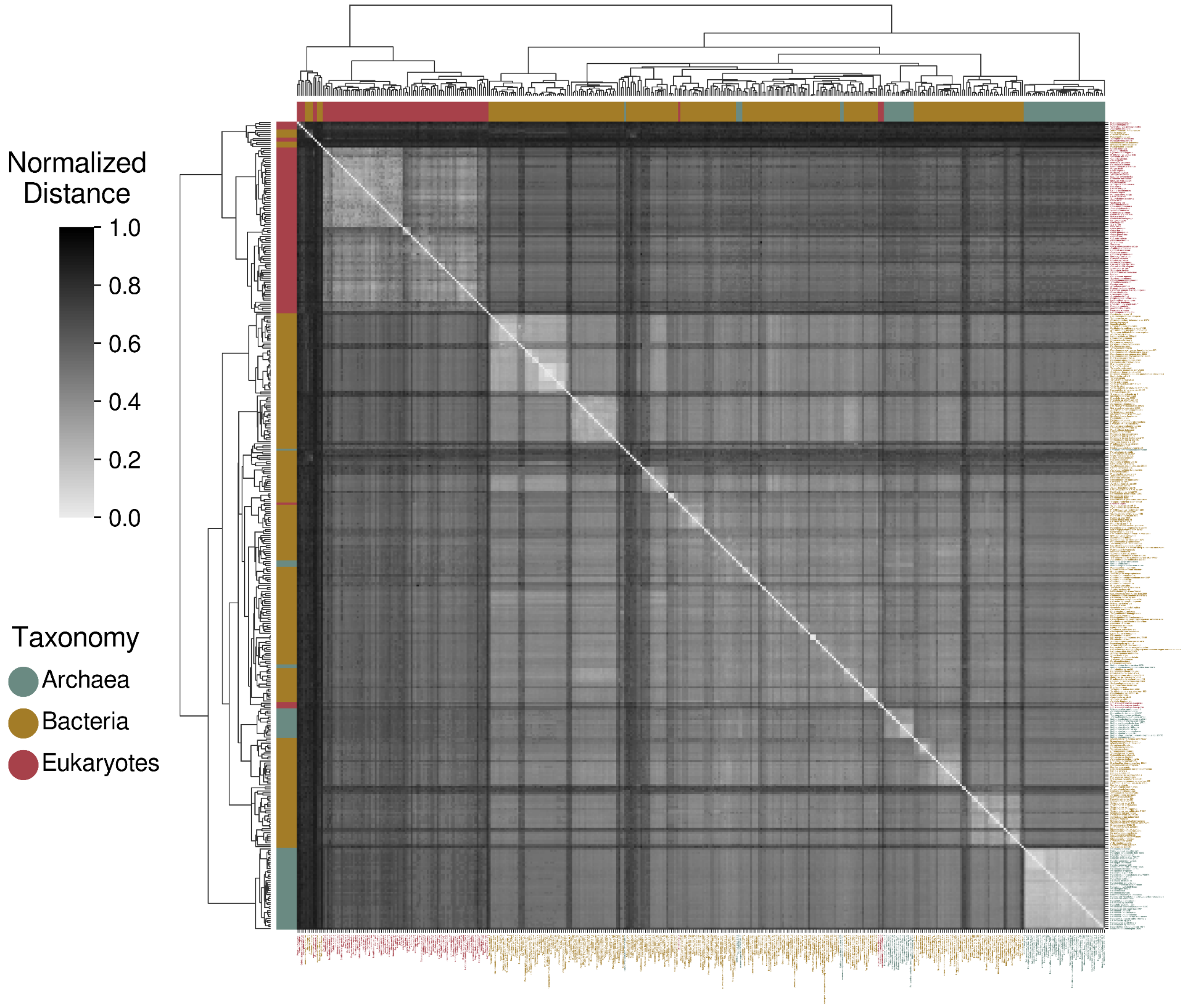


**Figure S2.4**: Phylogenetic clustergram derived from the comparison of heat shock protein 70kDa (HSP70) sequences. CD-HIT (Li & Godzik, 2006) and HMMER3 (Finn et al., 2011) were used to construct a consensus amino acid sequence for each species. The phylogenetic comparison was performed using the ClustalW tool (Sievers & Higgins, 2018).

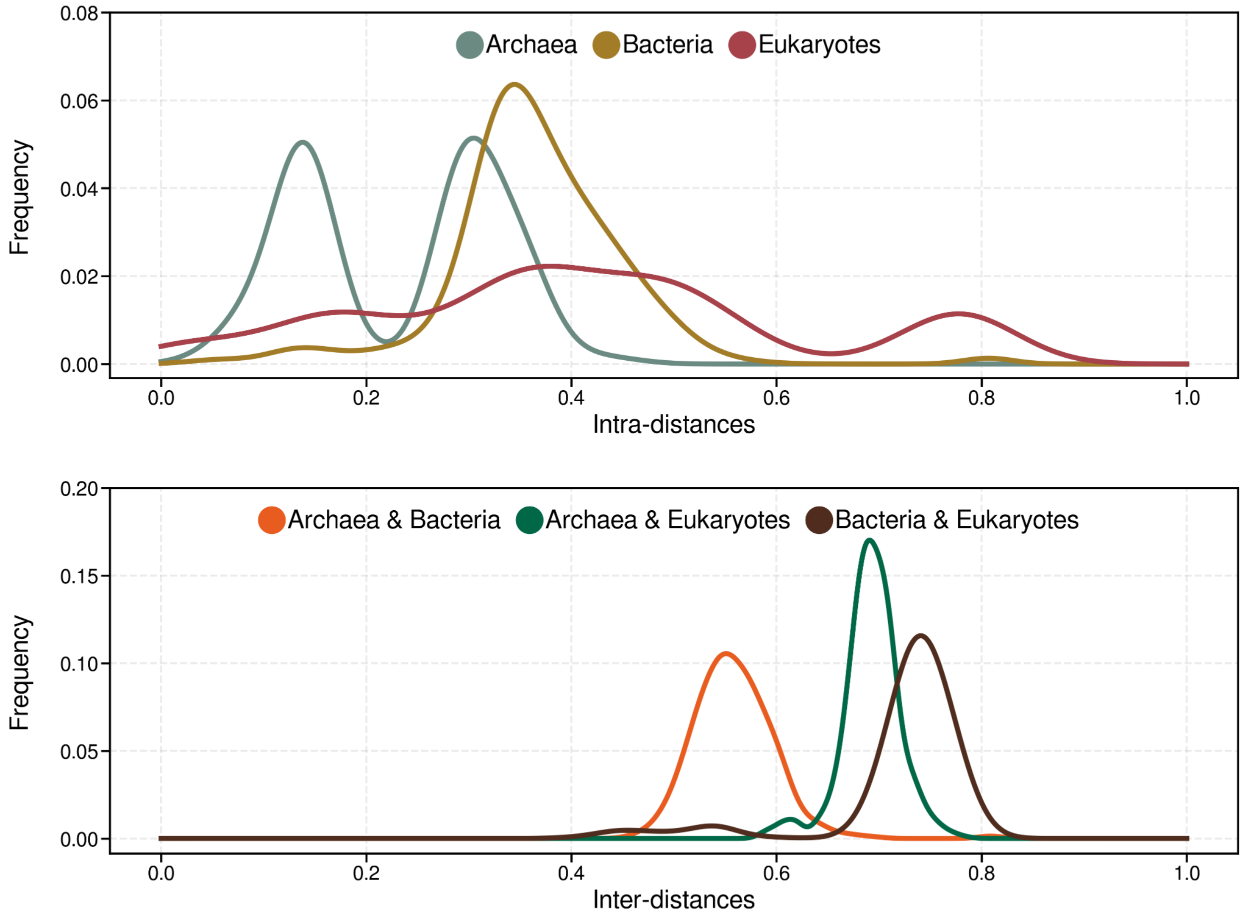


**Figure S2.5**: Distributions of pairwise distances of rRNA sequences in each taxonomic domain (intra-distances) and among the different domains (inter-distances).

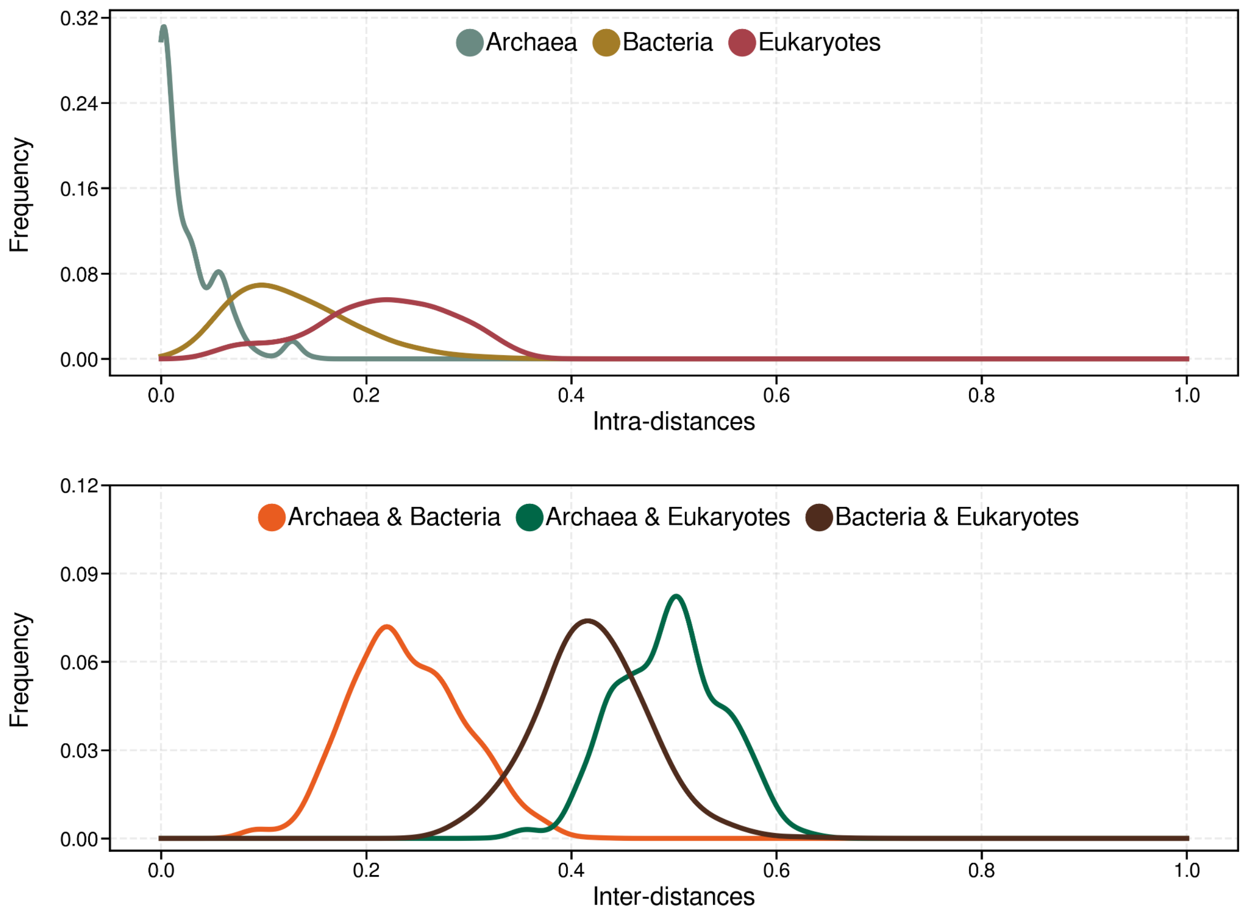


**Figure S2.6**: Distributions of pairwise PN-semantic distances in each taxonomic domain (intra-distances) and among the different domains (inter-distances).

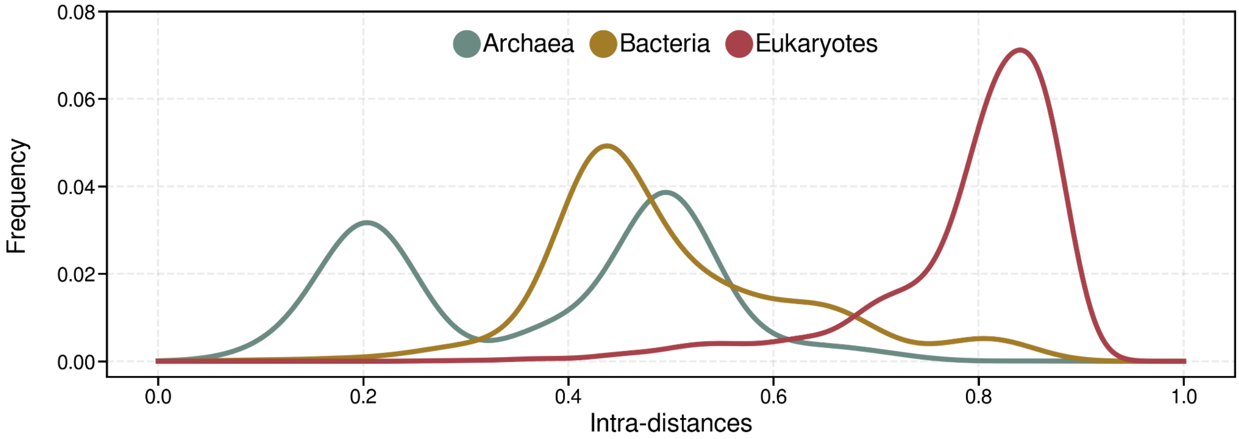


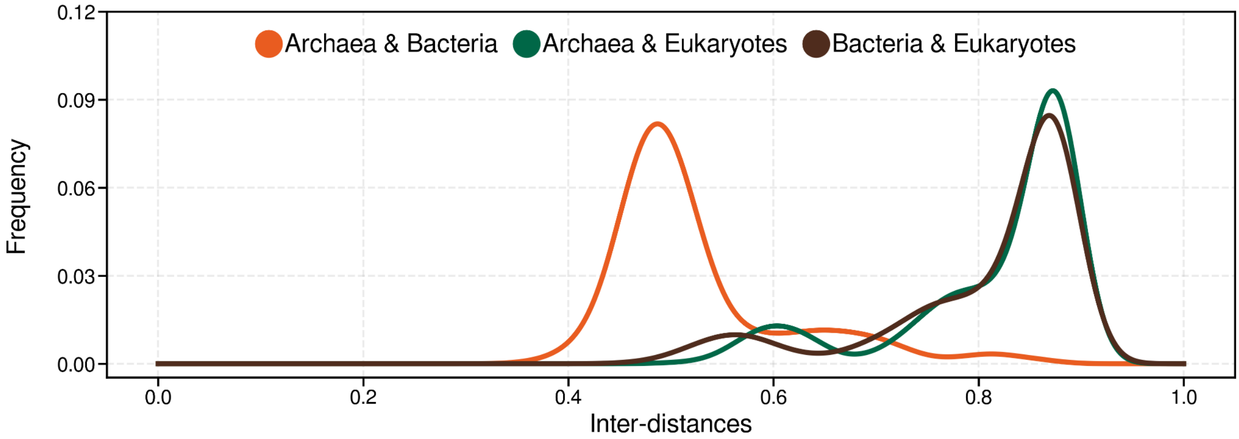


**Figure S2.7**: Distributions of pairwise distances of HSP40 sequences in each taxonomic domain (intra-distances) and among the different domains (inter-distances).

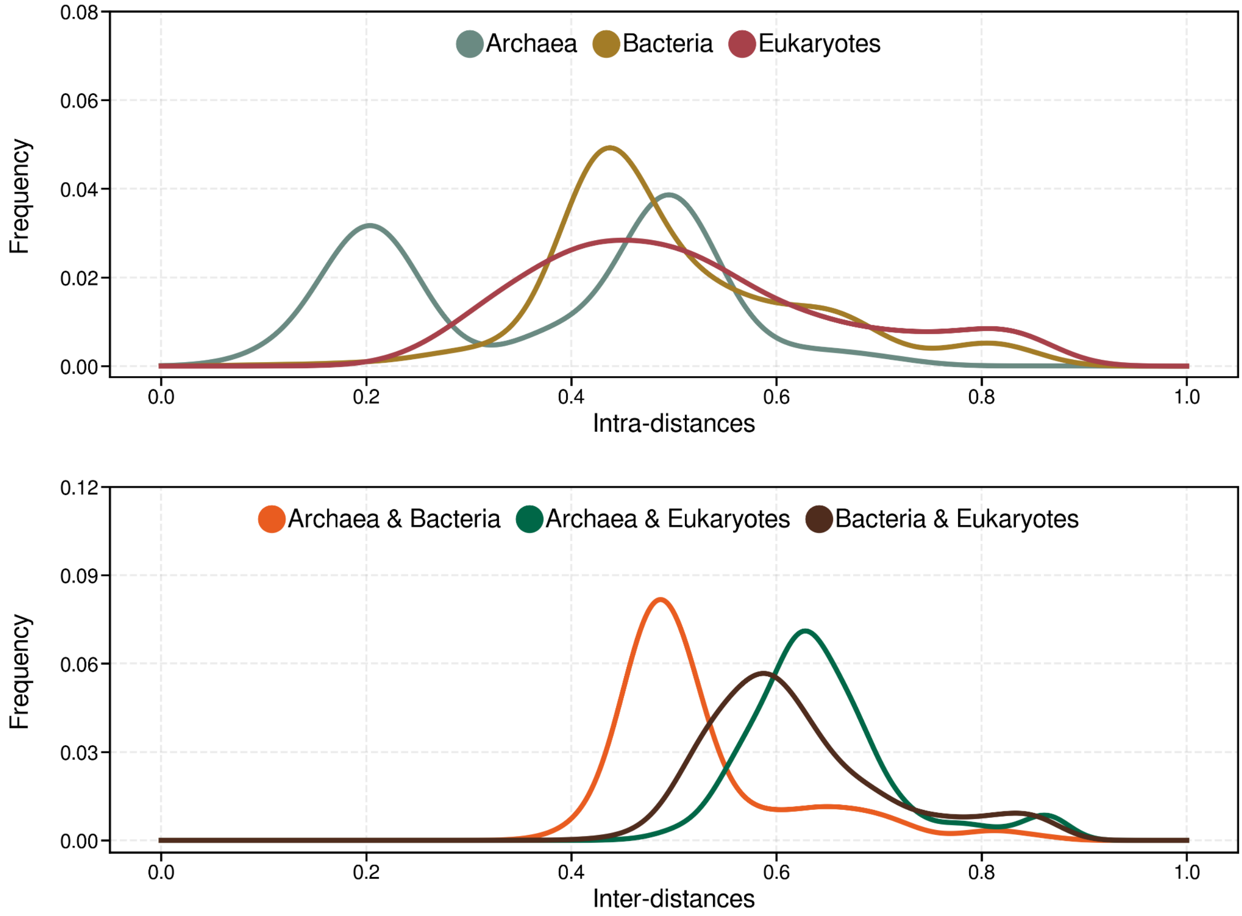


**Figure S2.8**: Distributions of pairwise distances of HSP70 sequences in each taxonomic domain (intra-distances) and among the different domains (inter-distances).

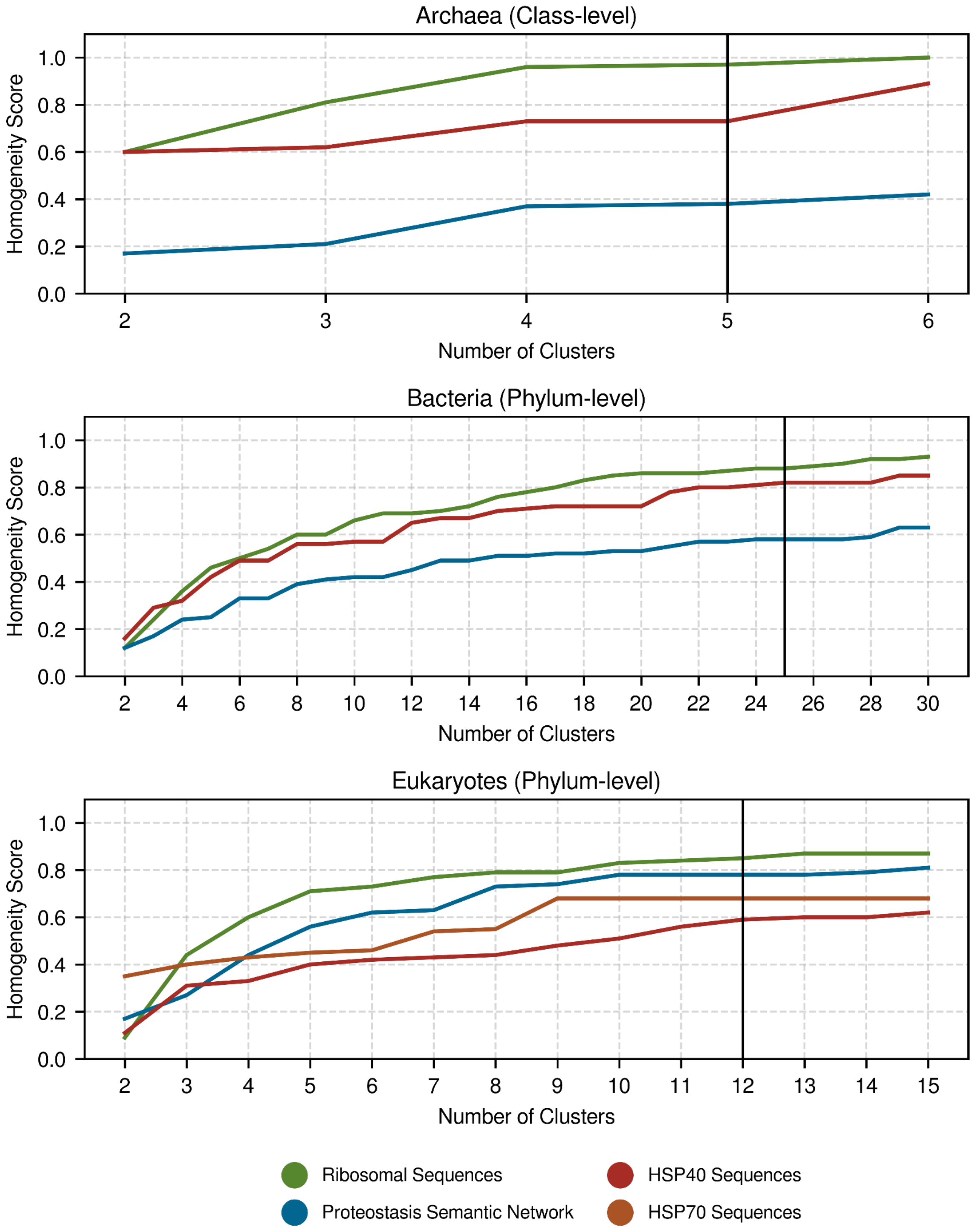


**Figure S2.9**: Evaluation of rRNA, HSP40, HSP70 and PN to separate effectively the species of the same domain into reference taxonomic sub-groups (Class-level for Archaea and Phylum-level for Bacteria and Eukaryotes). Different amounts of clusters were generated for each taxonomic category and the homogeneity score was calculated for each clustering outcome, based on the reference taxonomic classification of species. Vertical lines indicate the amount of reference taxonomic sub-groups.

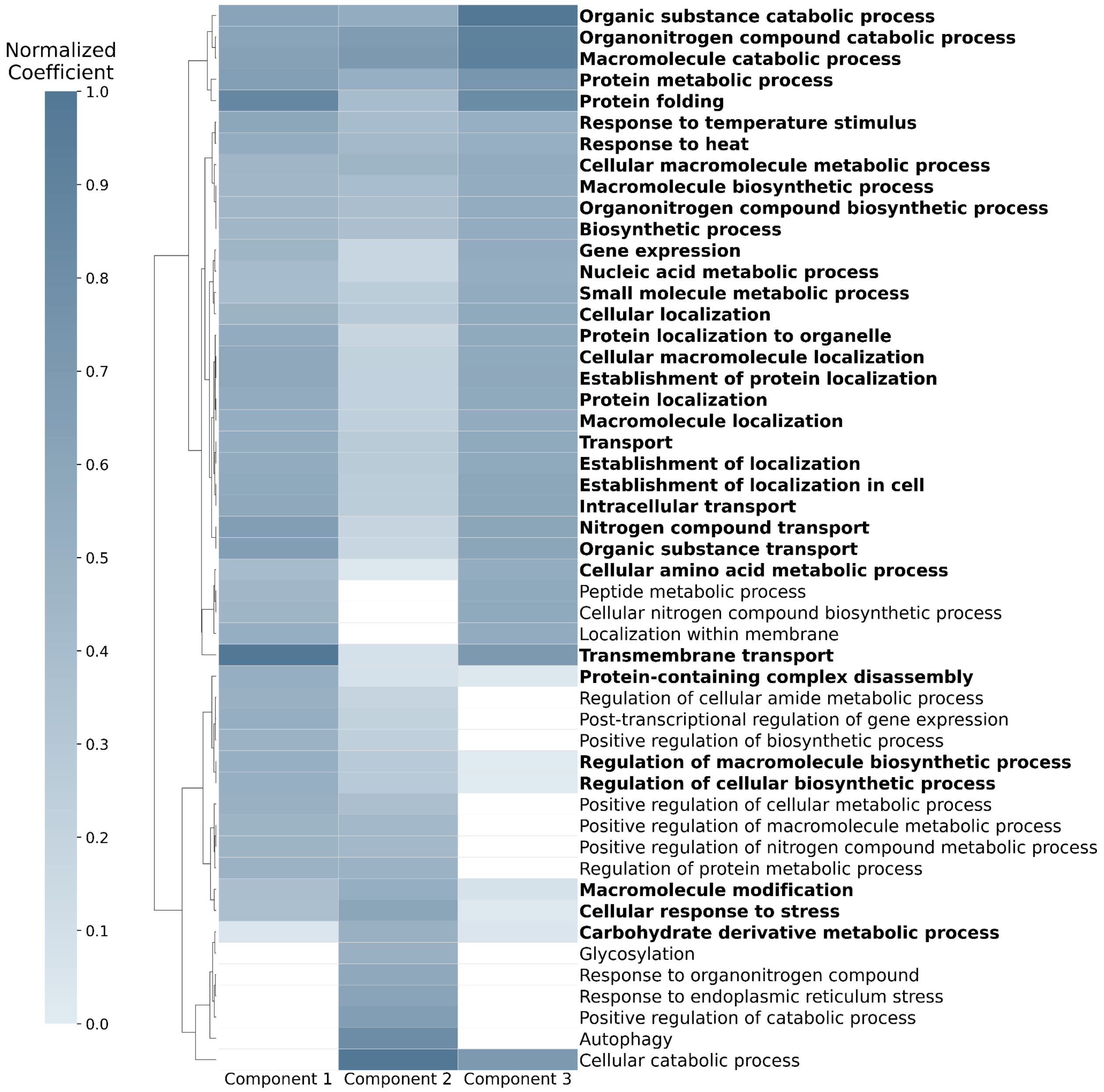


**Figure S2.10**: The features (components) matrix (n=3) obtained by applying non-negative matrix factorization to the association matrix between species and the PN-related semantic groups. The normalized coefficients indicate the contribution of each semantic group to the three NMF-based components. Bold semantic terms indicate the "conserved core" of PN profile across species.

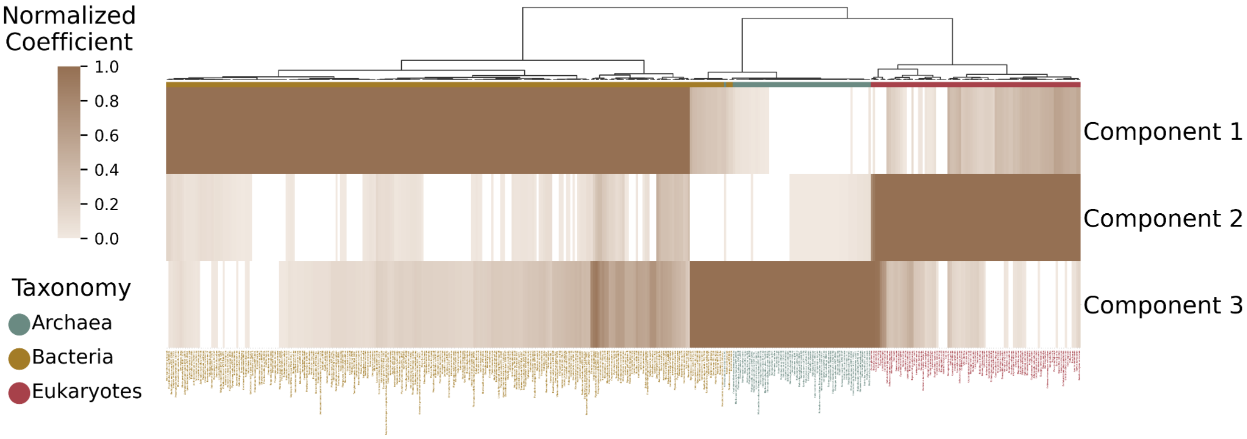


**Figure S2.11**: The coefficients matrix obtained by applying non-negative matrix factorization to the association matrix between species and the PN-related semantic groups. The normalized coefficients indicate the association between the species and NMF-based components.

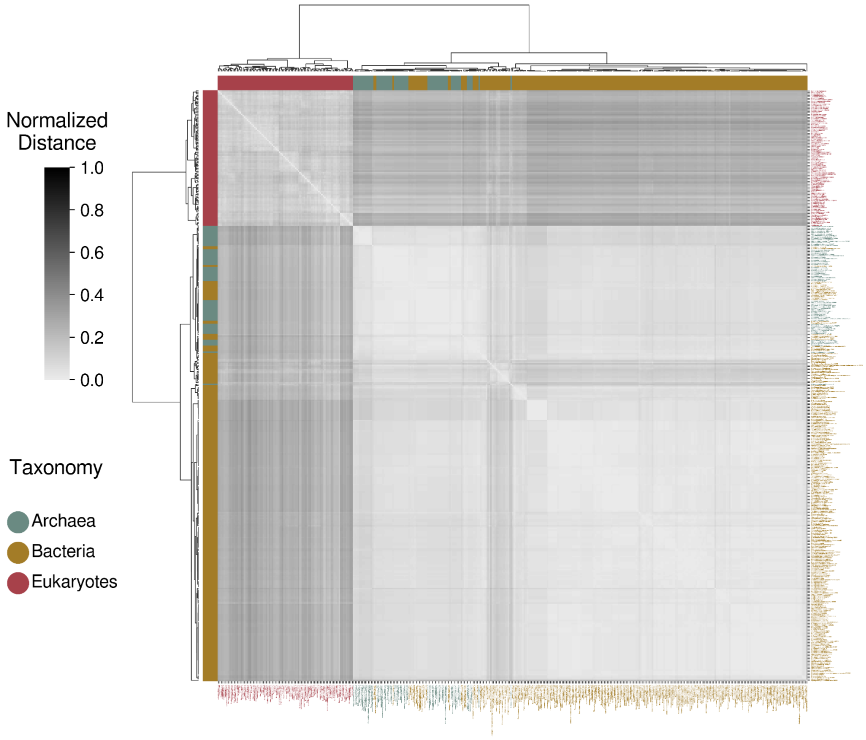


**Figure S2.12A**: Phylogenetic clustergram derived from the comparison of "ATP metabolic process" semantic networks.

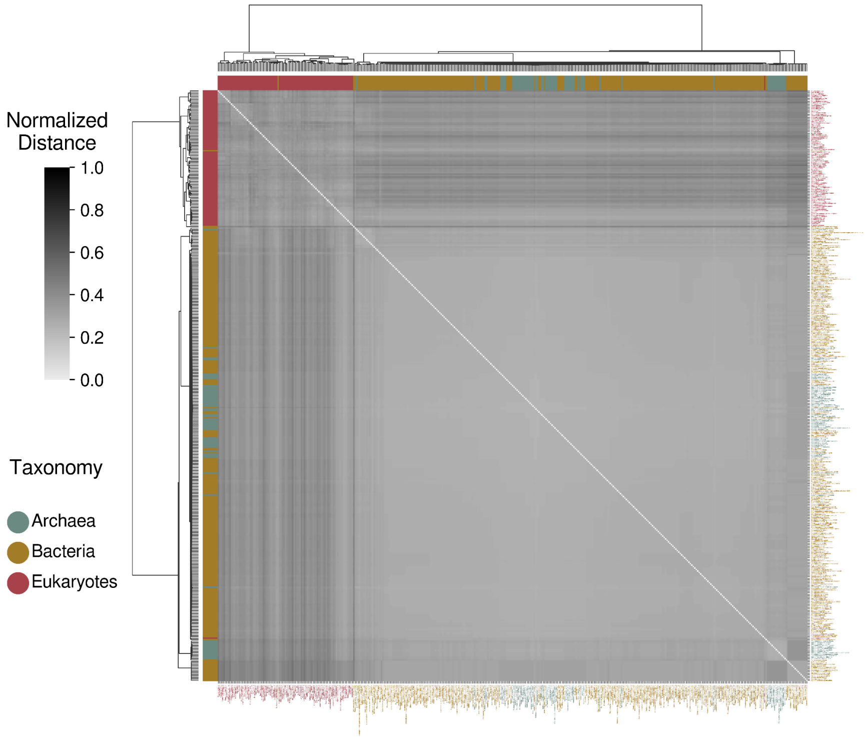


**Figure S2.12B**: Phylogenetic clustergram derived from the comparison of "ATP metabolic process" semantic networks, excluding terms associated with the obtained PN-related semantic groups.

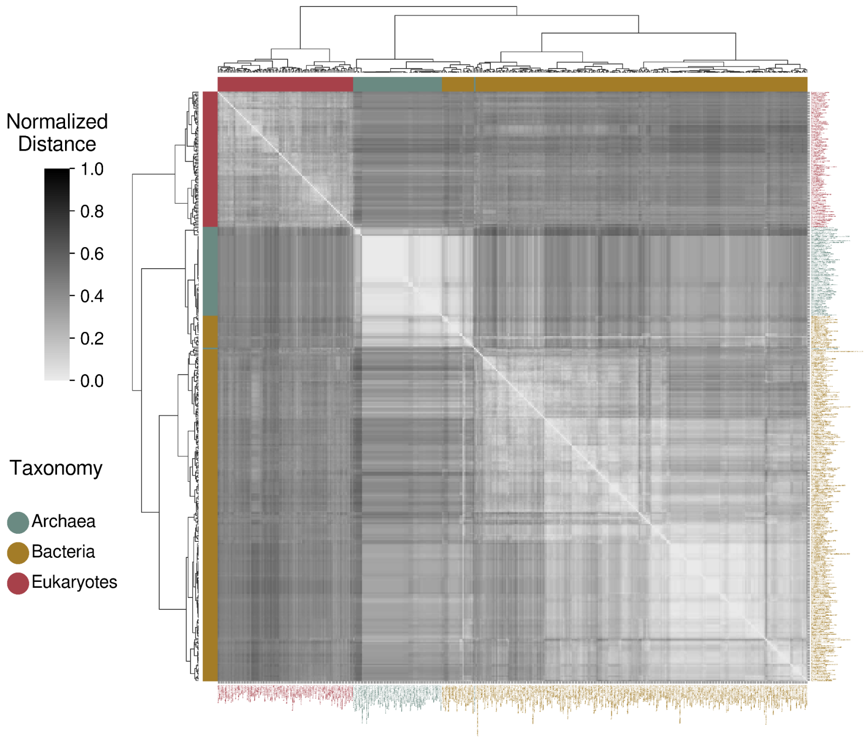


**Figure S2.13A**: Phylogenetic clustergram derived from the comparison of "cellular component assembly" semantic networks.

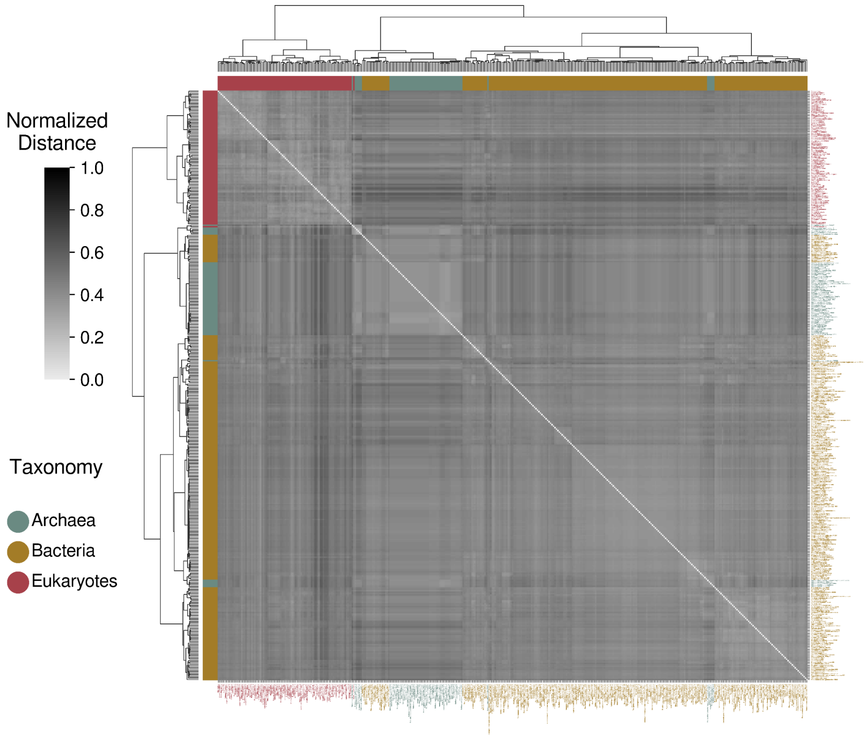


**Figure S2.13B**: Phylogenetic clustergram derived from the comparison of "cellular component assembly" semantic networks, excluding terms associated with the obtained PN-related semantic groups.

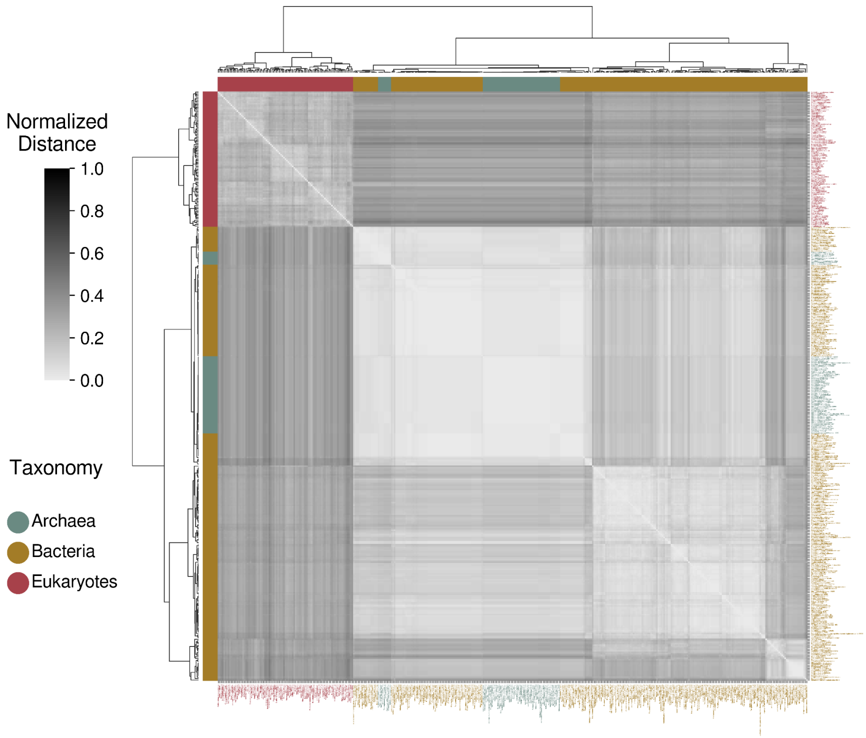


**Figure S2.14A**: Phylogenetic clustergram derived from the comparison of "cellular localization" semantic networks.

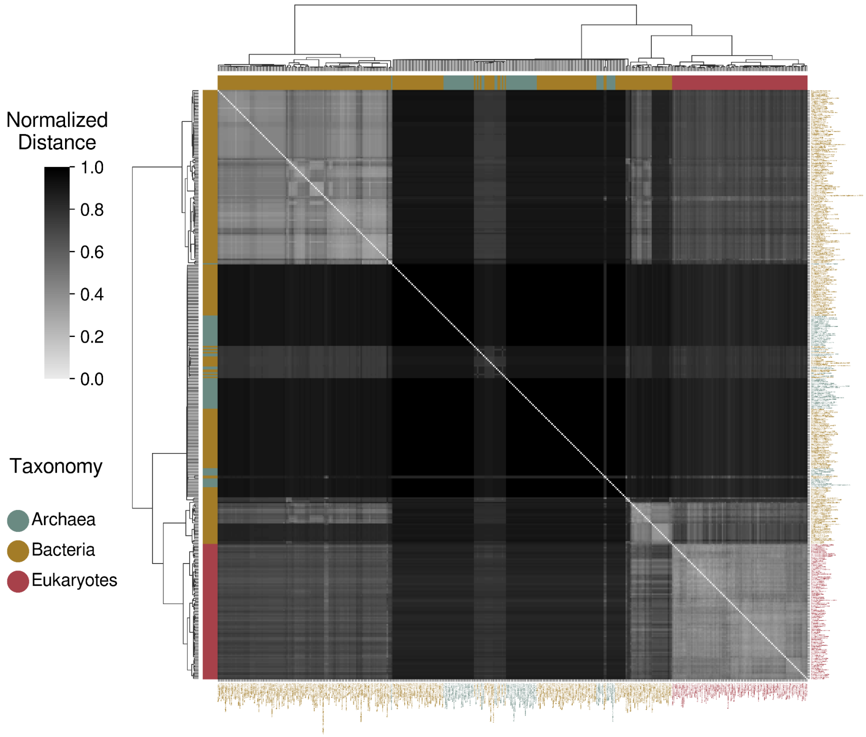


**Figure S2.14B**: Phylogenetic clustergram derived from the comparison of "cellular localization" semantic networks, excluding terms associated with the obtained PN-related semantic groups.

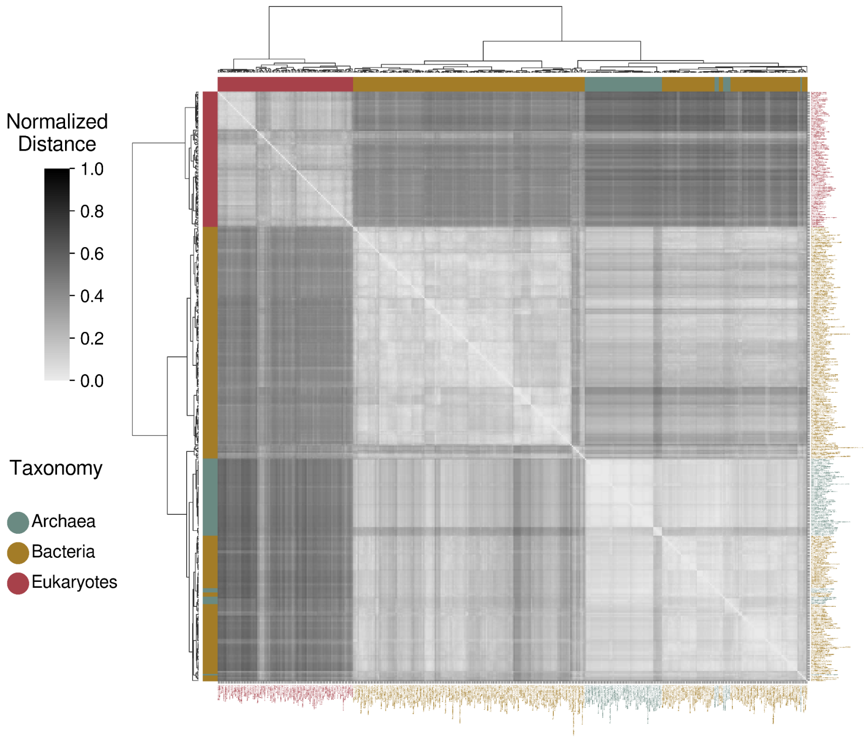


**Figure S2.15A**: Phylogenetic clustergram derived from the comparison of "cellular response to DNA damage stimulus" semantic networks.

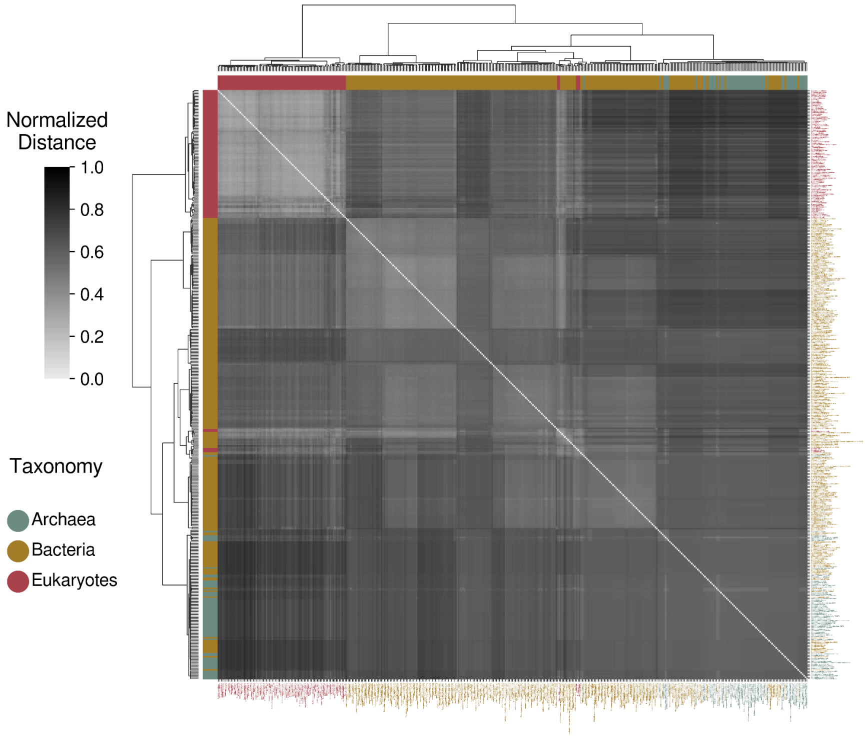


**Figure S2.15B**: Phylogenetic clustergram derived from the comparison of "cellular response to DNA damage stimulus" semantic networks, excluding terms associated with the obtained PN-related semantic groups.

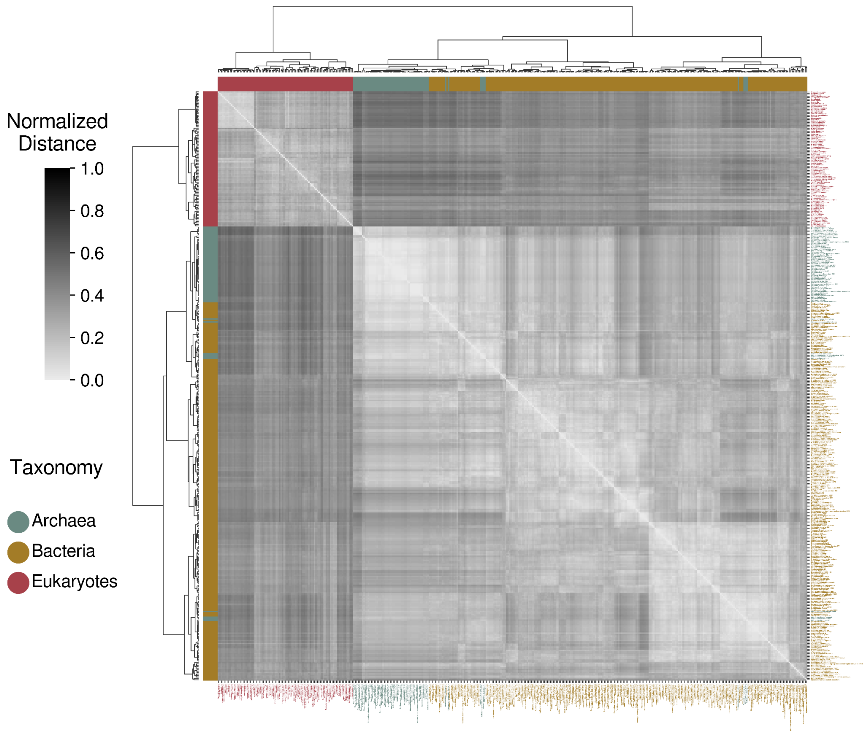


**Figure S2.16A**: Phylogenetic clustergram derived from the comparison of "cellular response to stress" semantic networks.

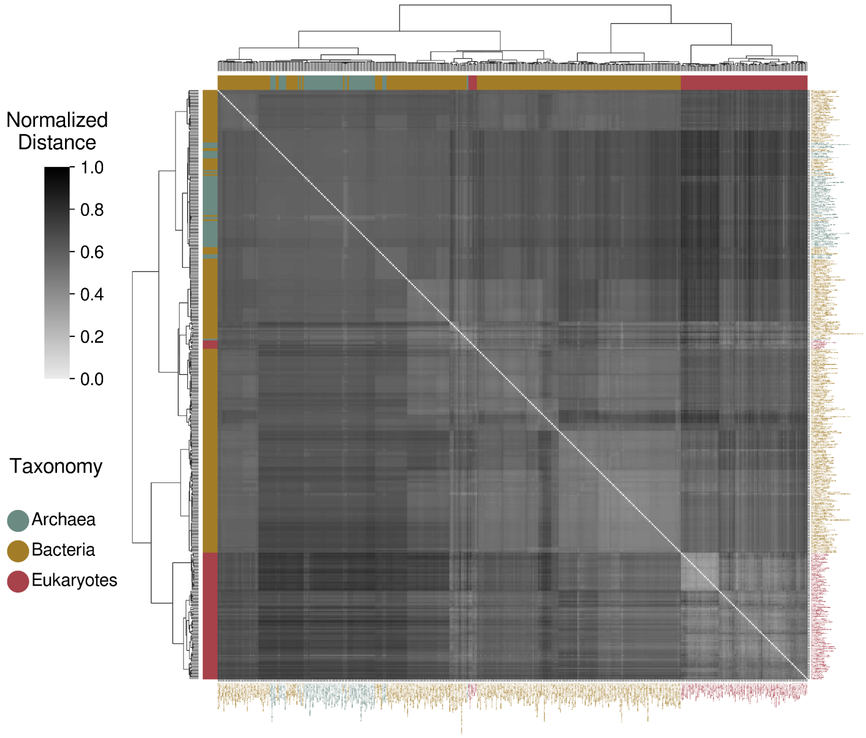


**Figure S2.16B**: Phylogenetic clustergram derived from the comparison of "cellular response to stress" semantic networks, excluding terms associated with the obtained PN-related semantic groups.

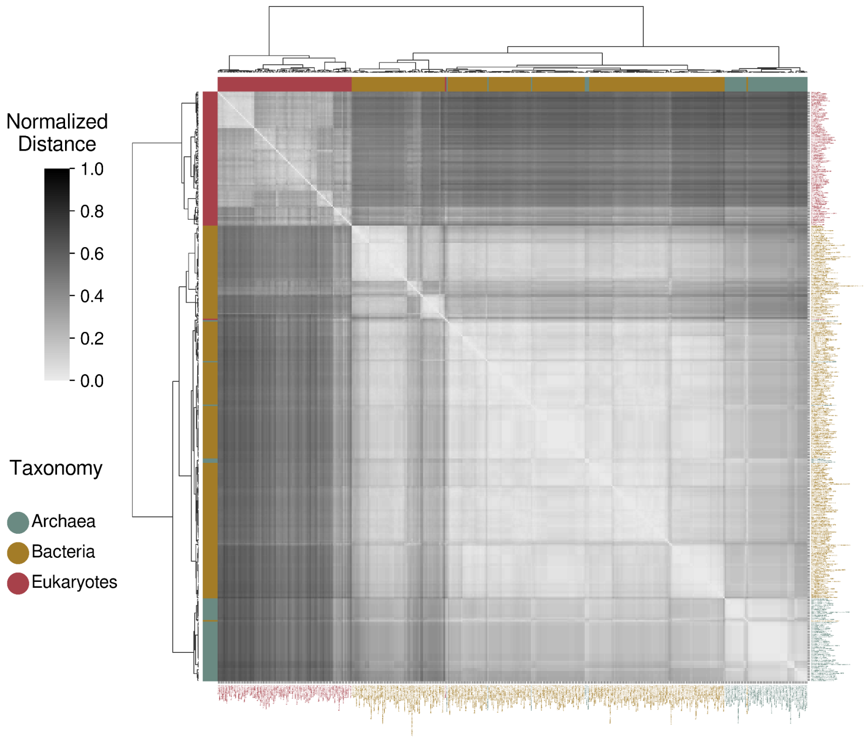


**Figure S2.17A**: Phylogenetic clustergram derived from the comparison of "DNA recombination" semantic networks.

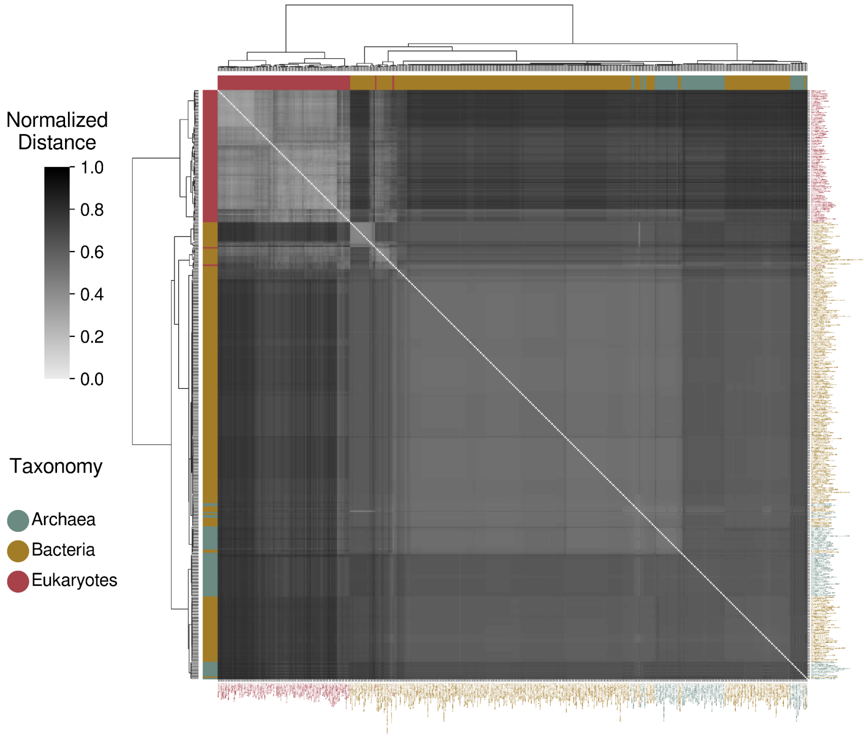


**Figure S2.17B**: Phylogenetic clustergram derived from the comparison of "DNA recombination" semantic networks, excluding terms associated with the obtained PN-related semantic groups.

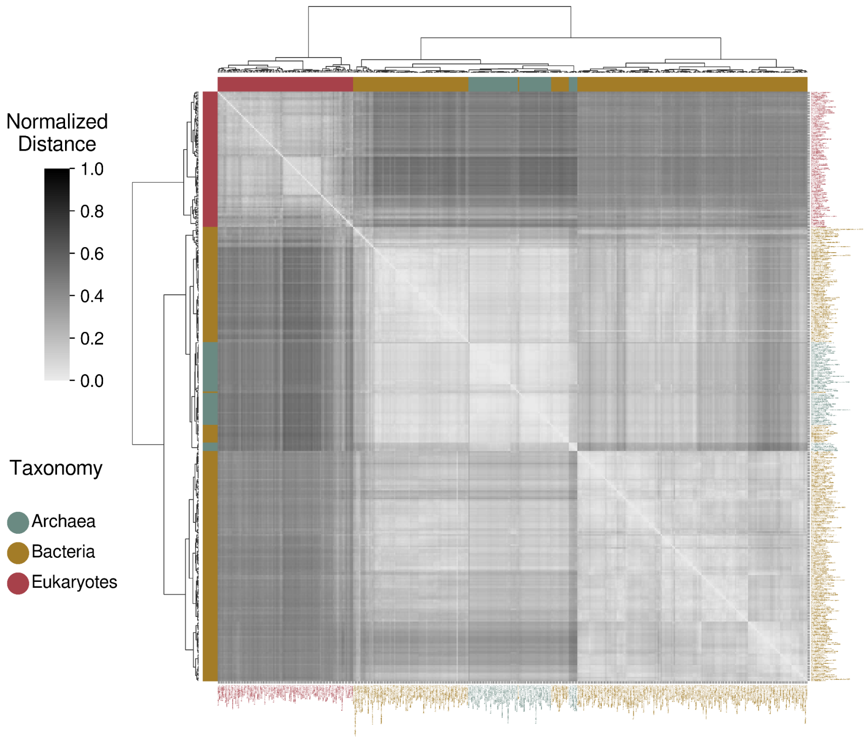


**Figure S2.18A**: Phylogenetic clustergram derived from the comparison of "DNA repair" semantic networks.

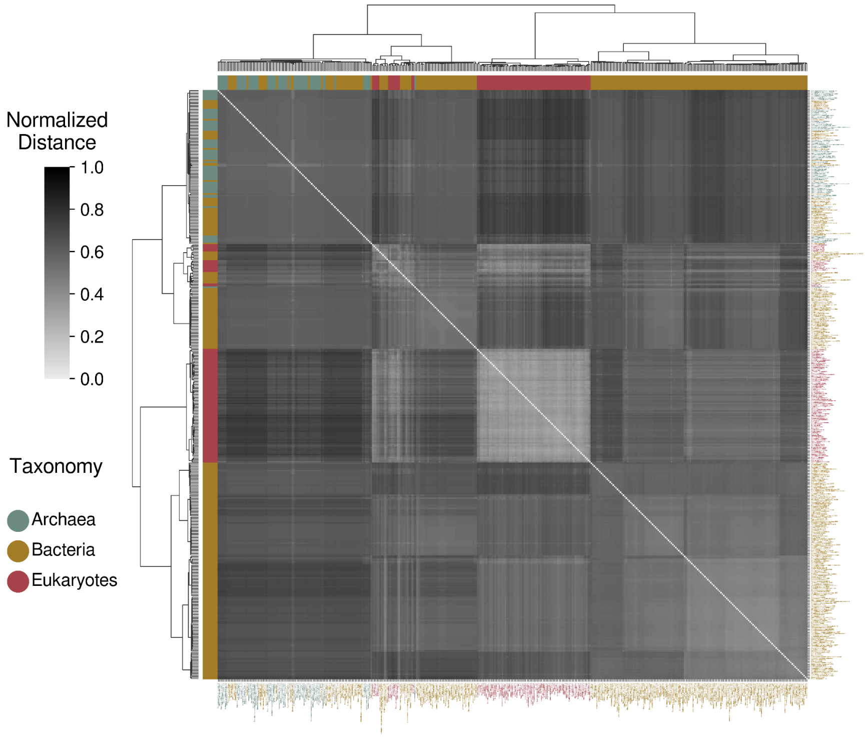


**Figure S2.18B**: Phylogenetic clustergram derived from the comparison of "DNA repair" semantic networks, excluding terms associated with the obtained PN-related semantic groups.

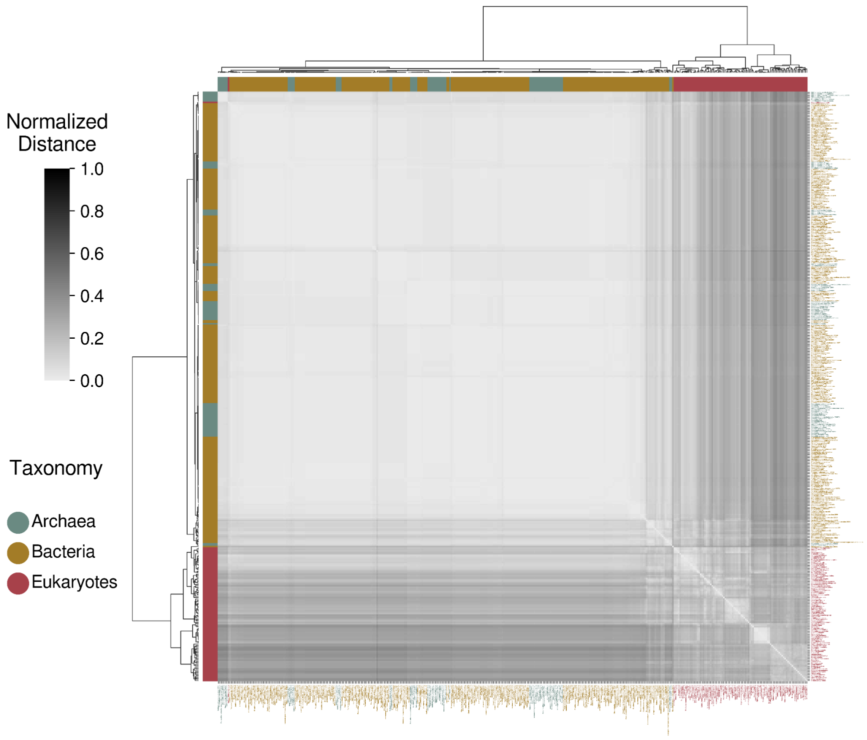


**Figure S2.19A**: Phylogenetic clustergram derived from the comparison of "glycolytic process" semantic networks.

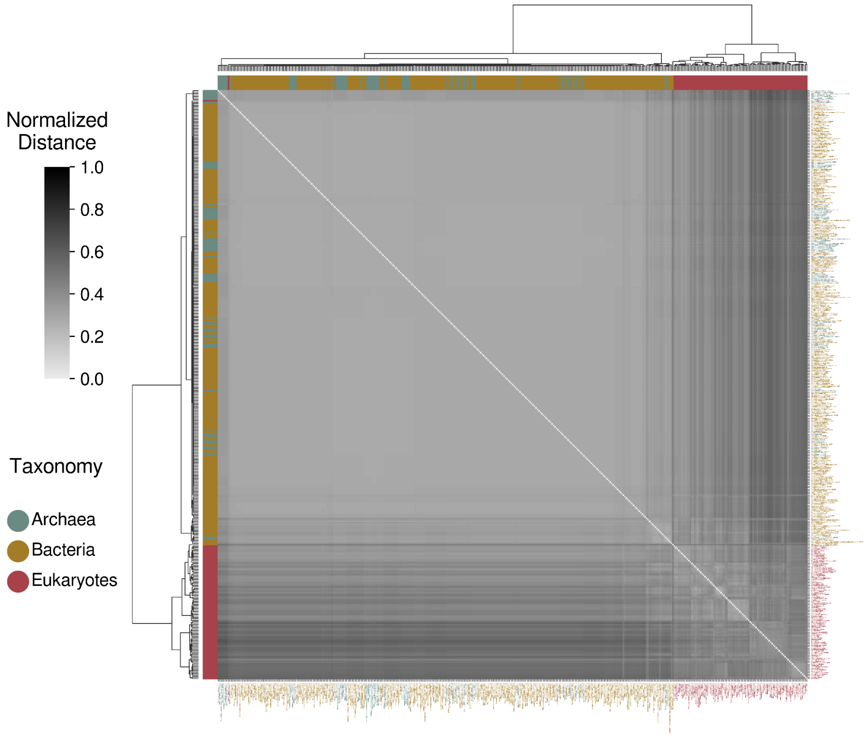


**Figure S2.19B**: Phylogenetic clustergram derived from the comparison of "glycolytic process" semantic networks, excluding terms associated with the obtained PN-related semantic groups.

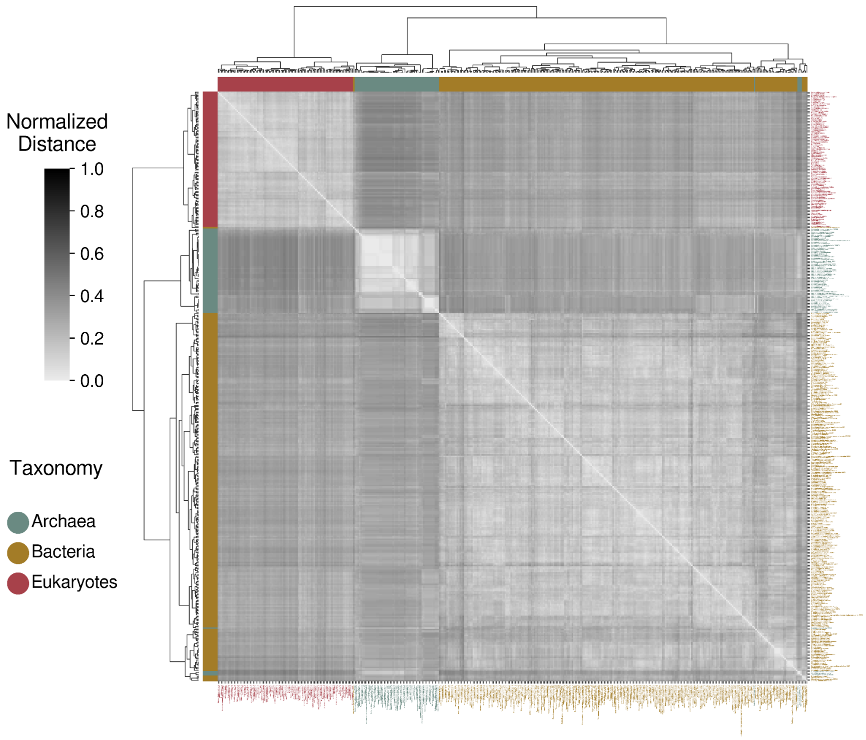


**Figure S2.20A**: Phylogenetic clustergram derived from the comparison of "lipid metabolic process" semantic networks.

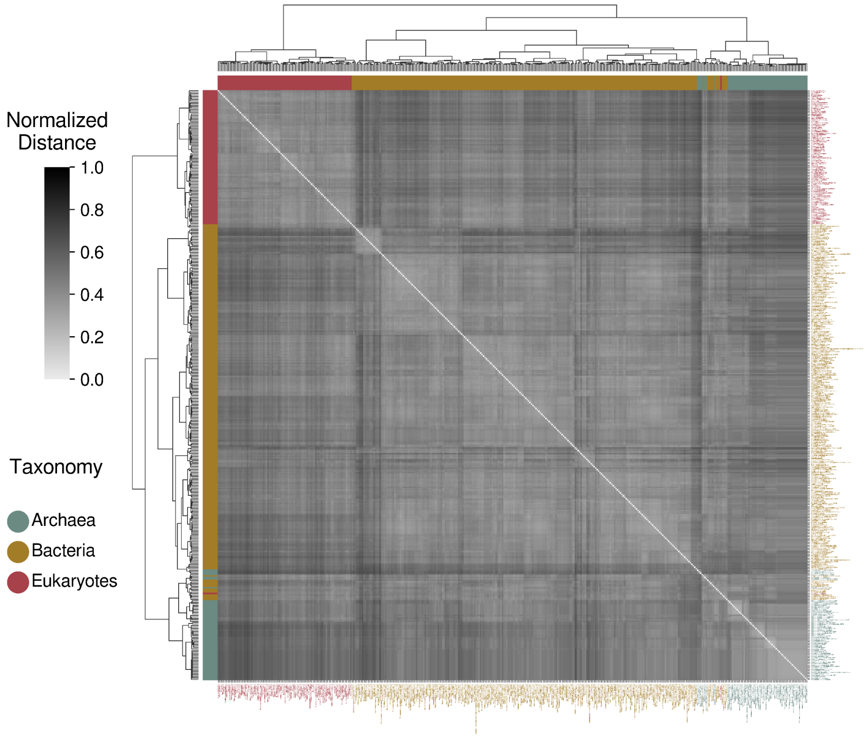


**Figure S2.20B**: Phylogenetic clustergram derived from the comparison of "lipid metabolic process" semantic networks, excluding terms associated with the obtained PN-related semantic groups.

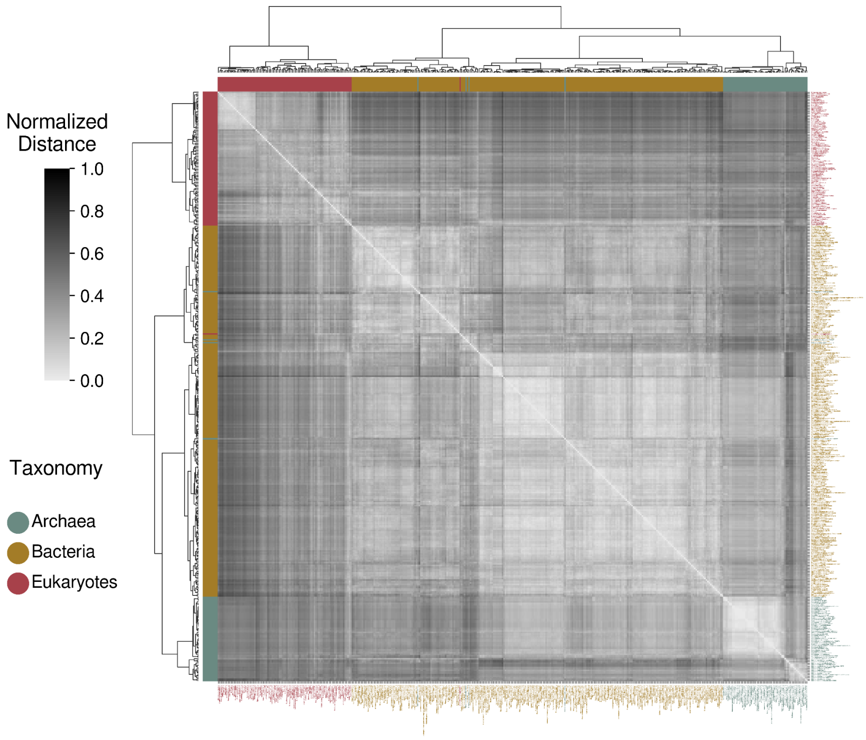


**Figure S2.21A**: Phylogenetic clustergram derived from the comparison of "methylation" semantic networks.

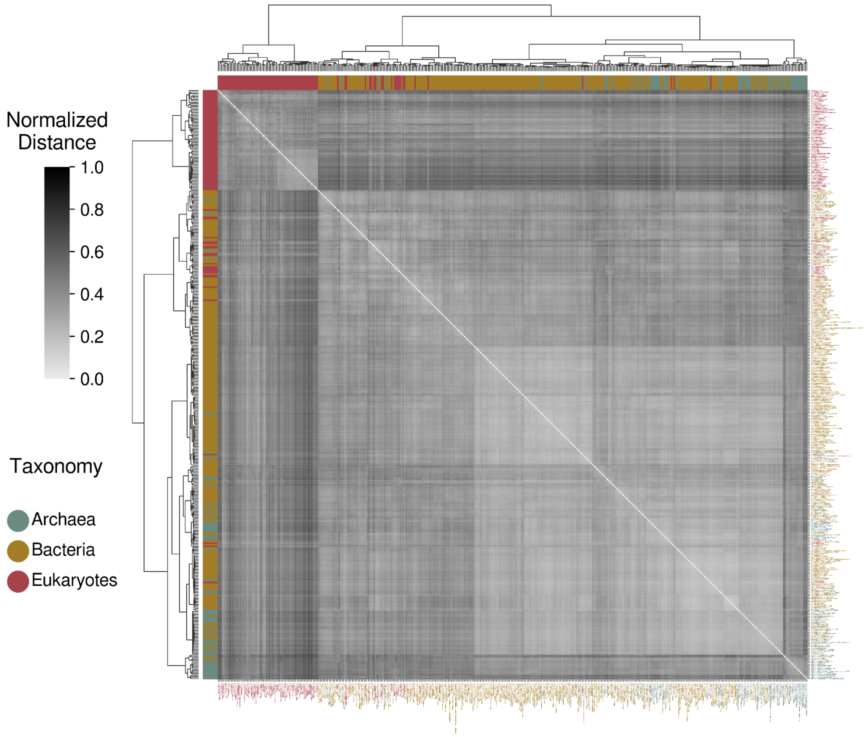


**Figure S2.21B**: Phylogenetic clustergram derived from the comparison of "methylation" semantic networks, excluding terms associated with the obtained PN-related semantic groups.

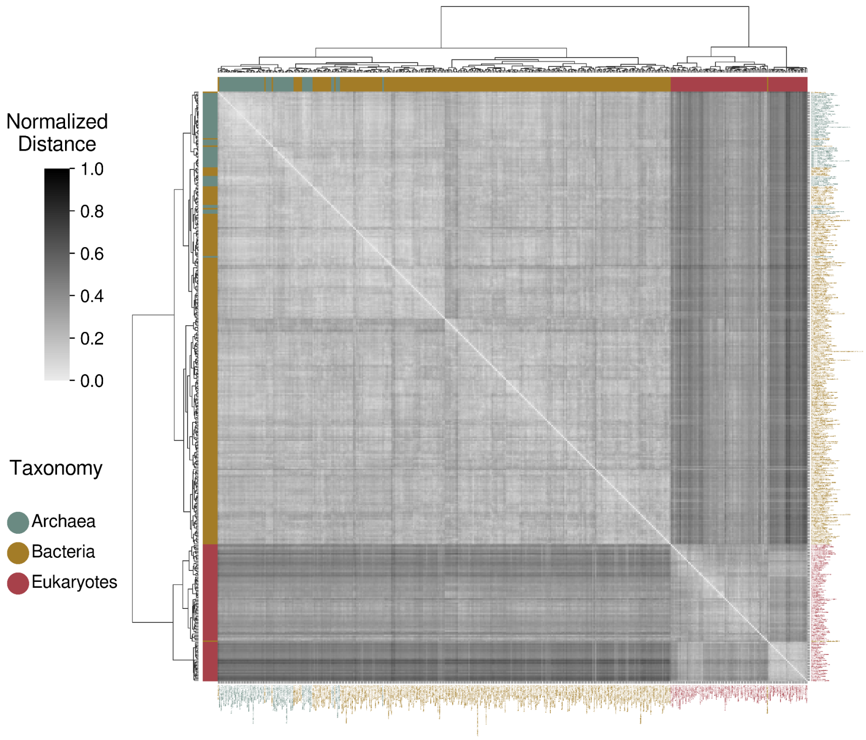


**Figure S2.22A**: Phylogenetic clustergram derived from the comparison of "phosphorylation" semantic networks.

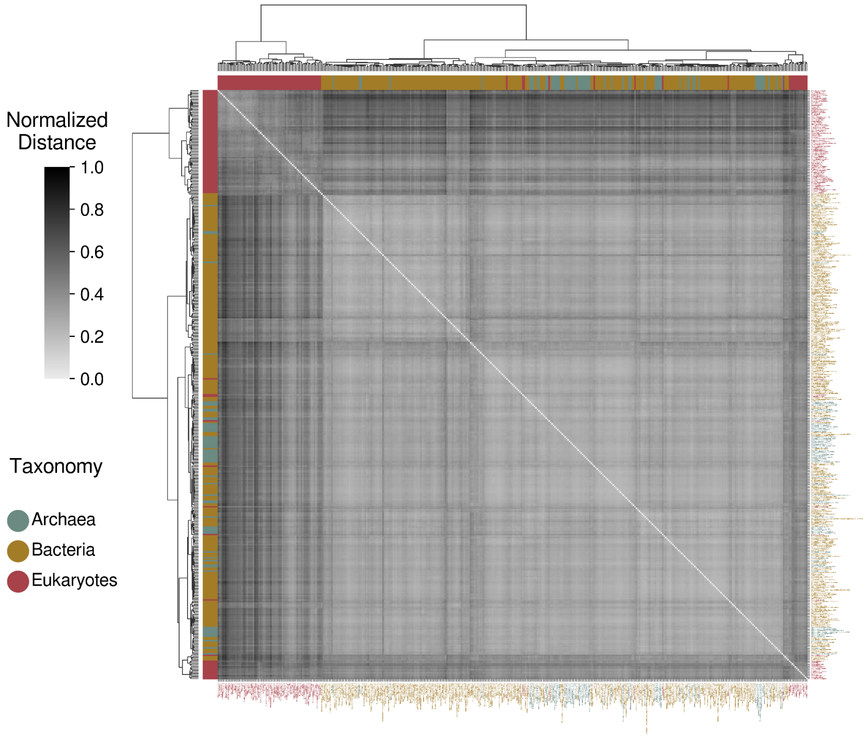


**Figure S2.22B**: Phylogenetic clustergram derived from the comparison of "phosphorylation" semantic networks, excluding terms associated with the obtained PN-related semantic groups.

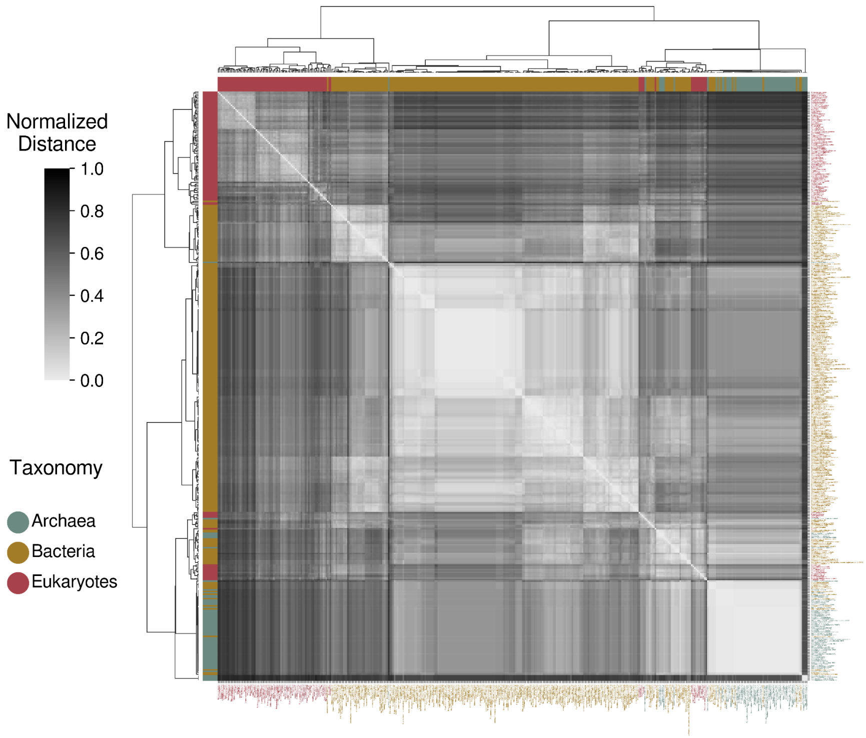


**Figure S2.23A**: Phylogenetic clustergram derived from the comparison of "protein folding" semantic networks.

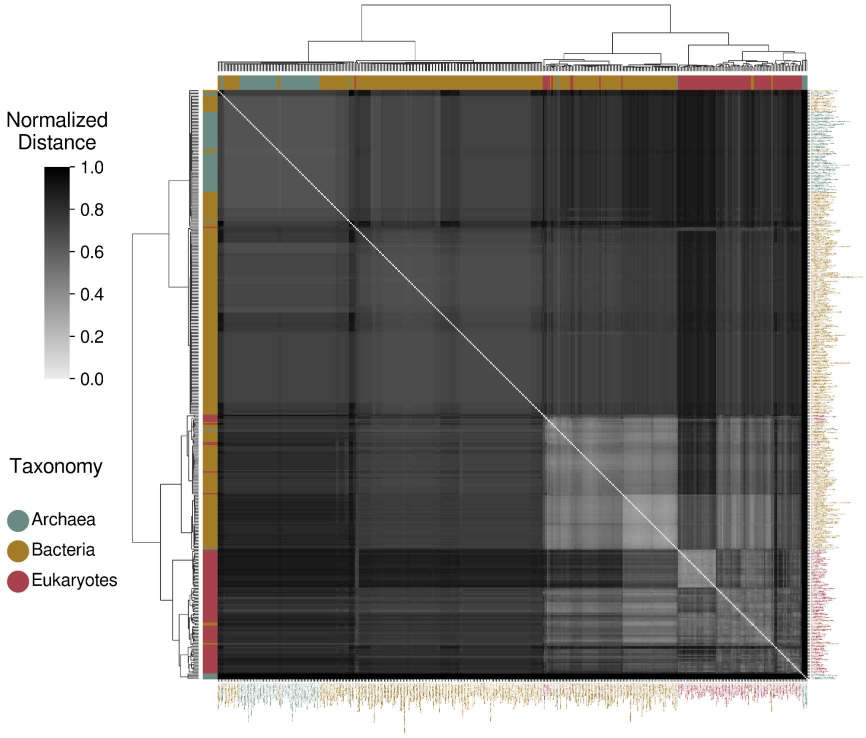


**Figure S2.23B**: Phylogenetic clustergram derived from the comparison of "protein folding" semantic networks, excluding terms associated with the obtained PN-related semantic groups.

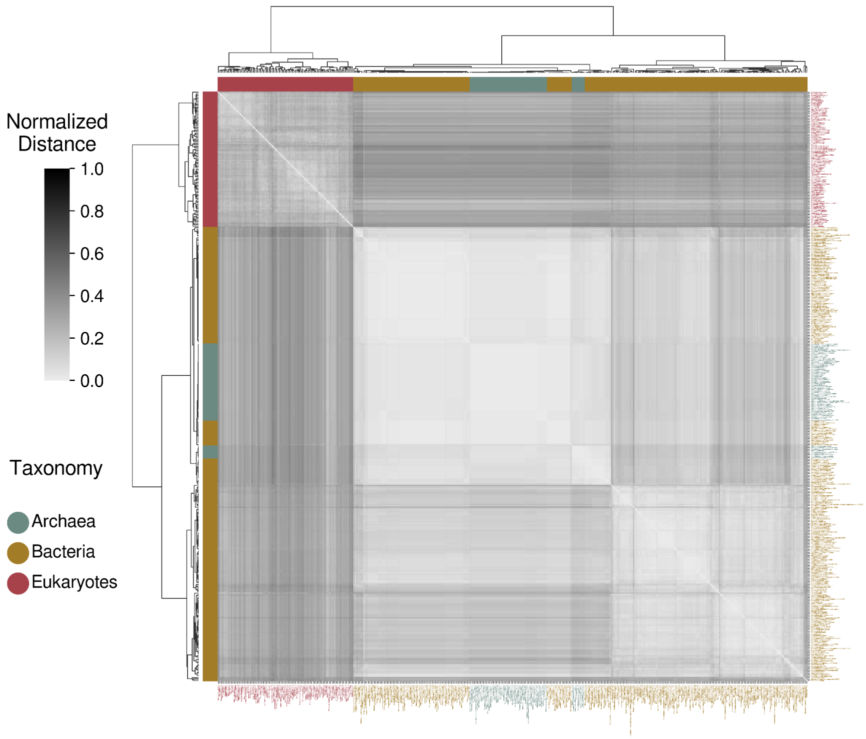


**Figure S2.24A**: Phylogenetic clustergram derived from the comparison of “protein localization” semantic networks.

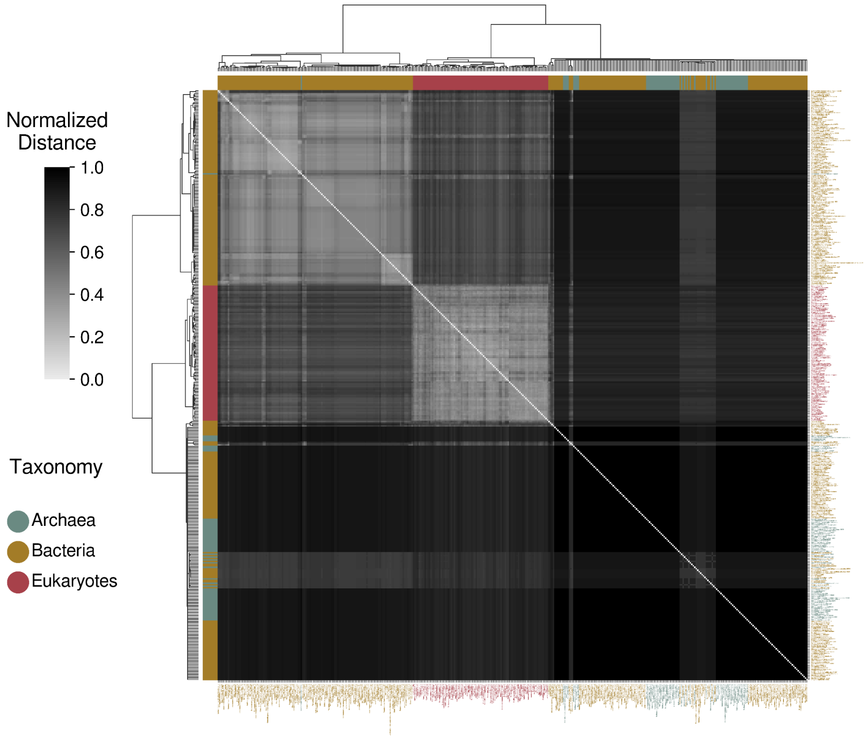


**Figure S2.24B**: Phylogenetic clustergram derived from the comparison of “protein localization” semantic networks, excluding terms associated with the obtained PN-related semantic groups.

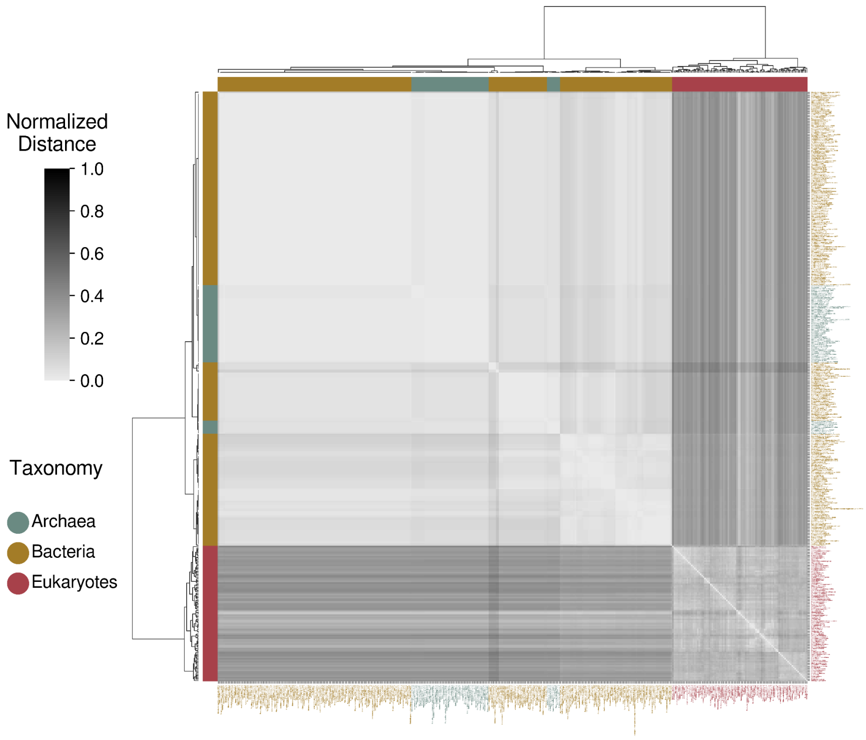


**Figure S2.25A**: Phylogenetic clustergram derived from the comparison of "protein targeting" semantic networks.

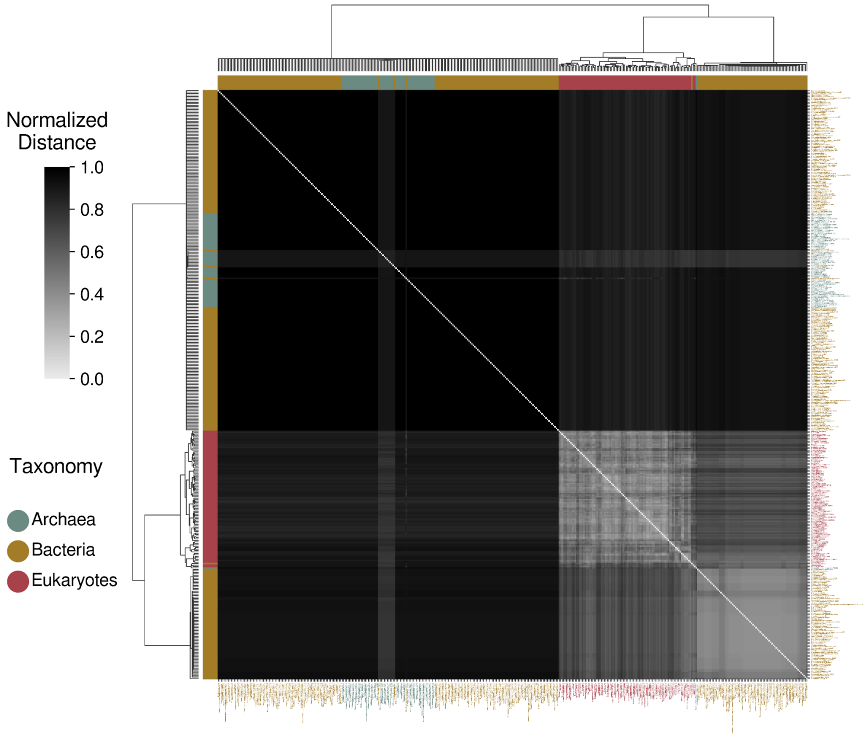


**Figure S2.25B**: Phylogenetic clustergram derived from the comparison of "protein targeting" semantic networks, excluding terms associated with the obtained PN-related semantic groups.

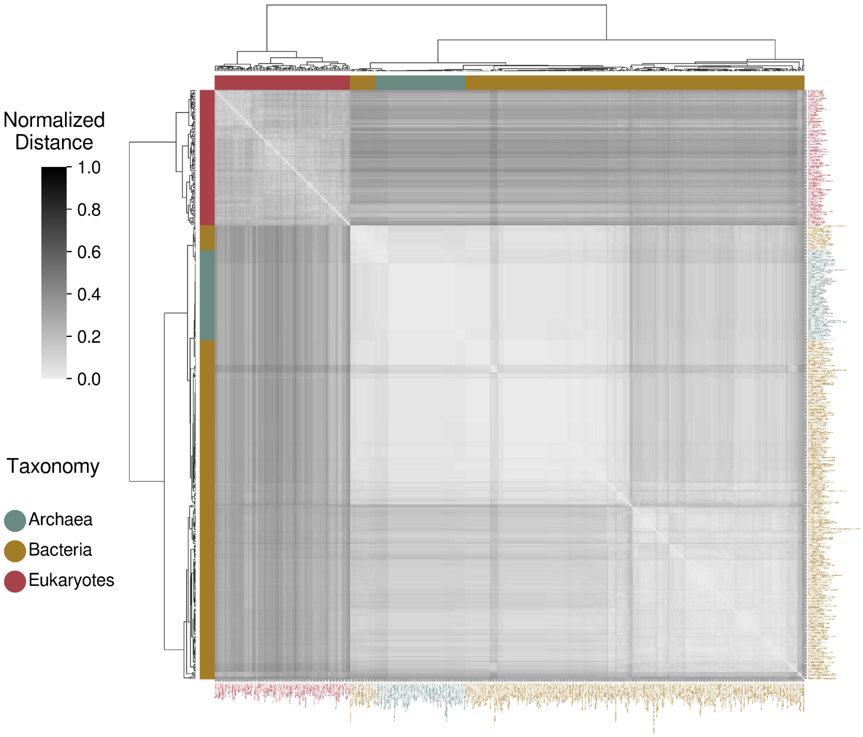


**Figure S2.26A**: Phylogenetic clustergram derived from the comparison of "protein transport" semantic networks.

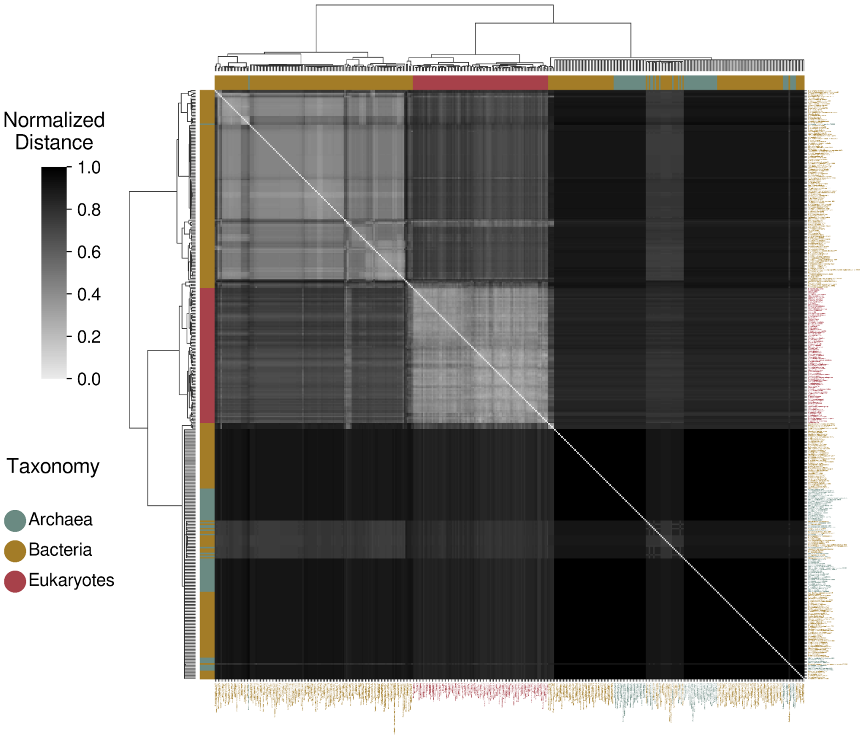


**Figure S2.26B**: Phylogenetic clustergram derived from the comparison of "protein transport" semantic networks, excluding terms associated with the obtained PN-related semantic groups.

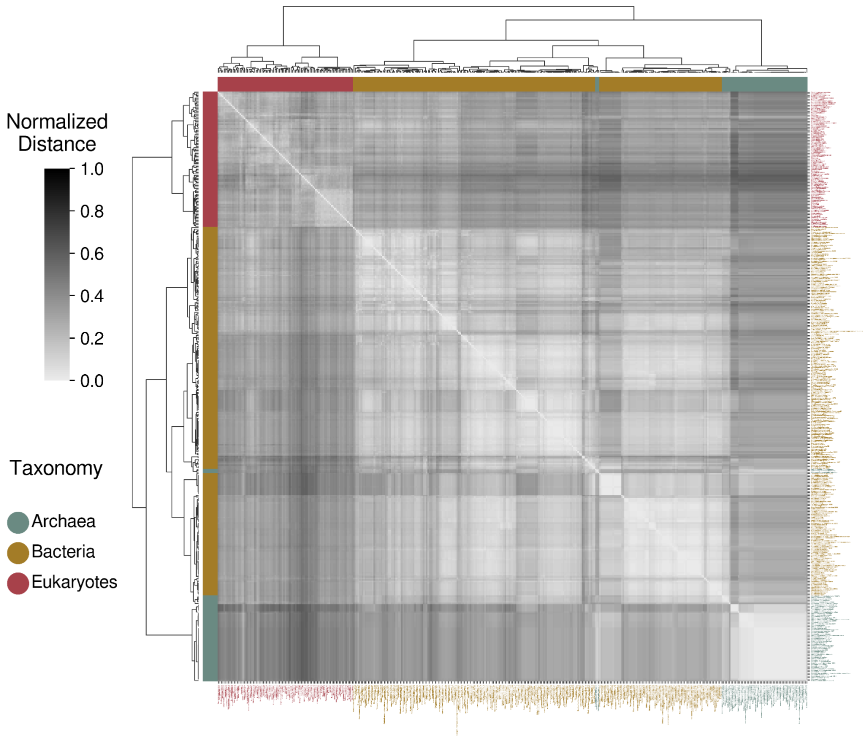


**Figure S2.27A**: Phylogenetic clustergram derived from the comparison of "regulation of DNA-templated transcription" semantic networks.

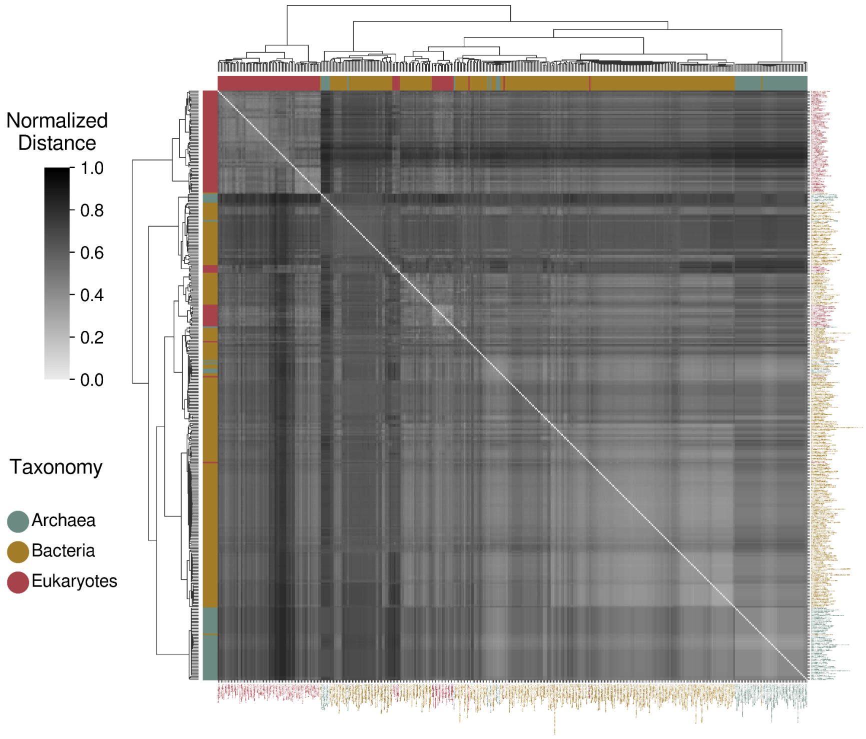


**Figure S2.27B**: Phylogenetic clustergram derived from the comparison of "regulation of DNA-templated transcription" semantic networks, excluding terms associated with the obtained PN-related semantic groups.

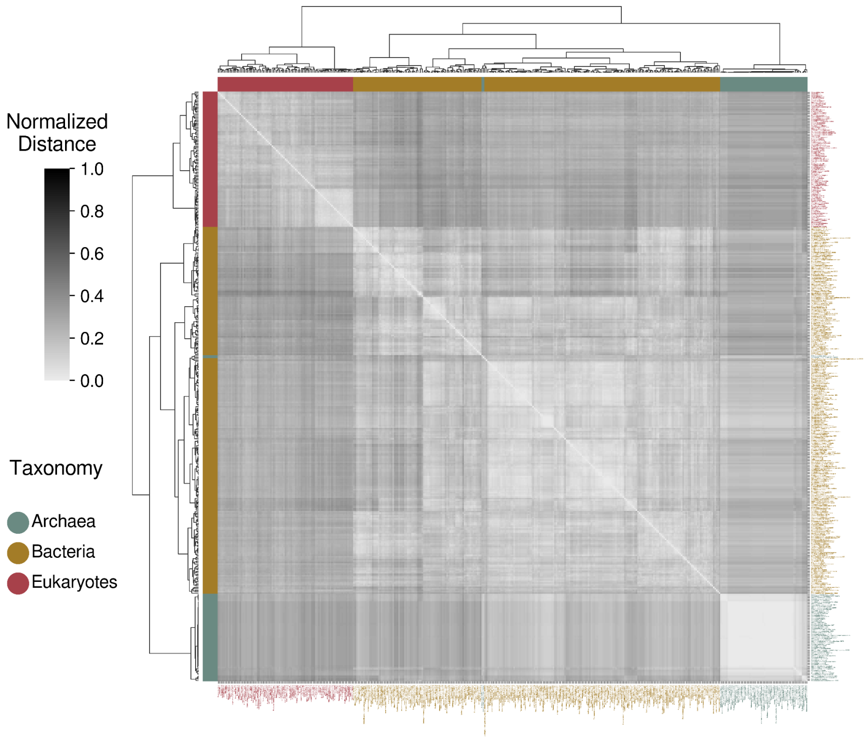


**Figure S2.28A**: Phylogenetic clustergram derived from the comparison of “regulation of macromolecule metabolic process” semantic networks.

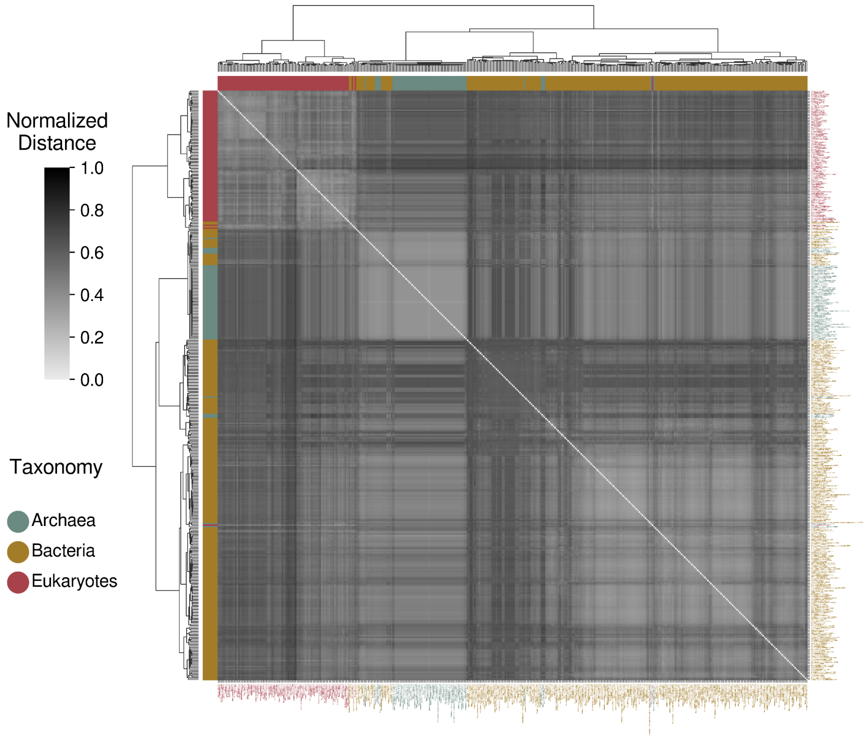


**Figure S2.28B**: Phylogenetic clustergram derived from the comparison of “regulation of macromolecule metabolic process” semantic networks, excluding terms associated with the obtained PN-related semantic groups.

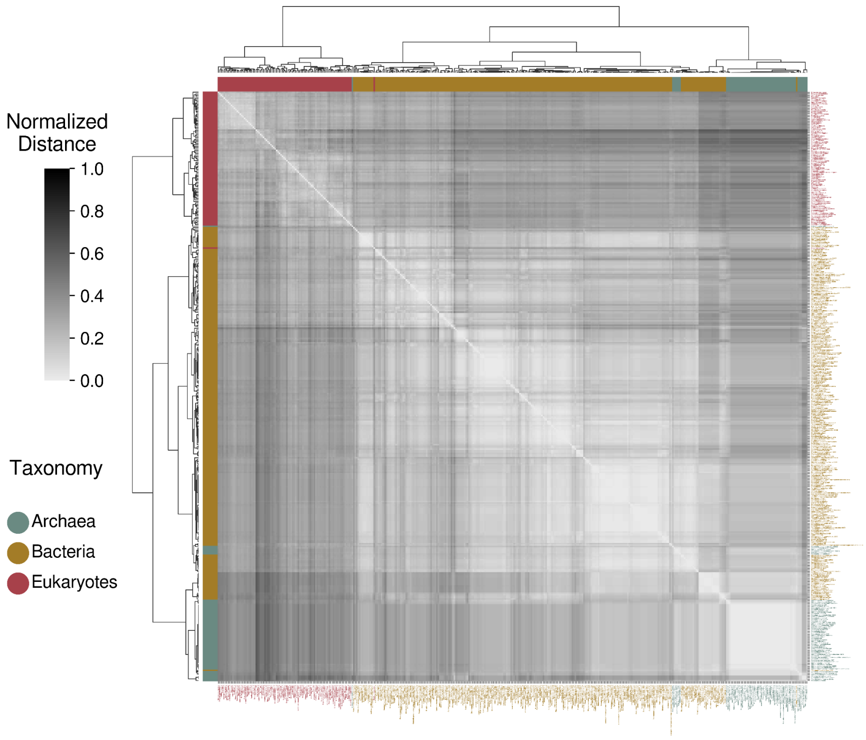


**Figure S2.29A**: Phylogenetic clustergram derived from the comparison of "regulation of RNA metabolic process" semantic networks.

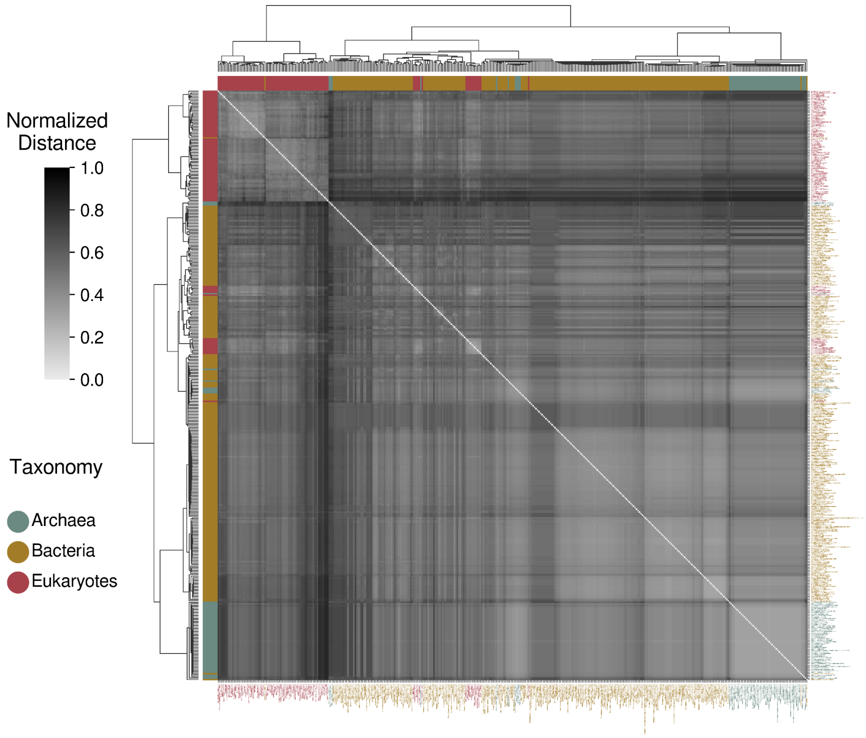


**Figure S2.29B**: Phylogenetic clustergram derived from the comparison of "regulation of RNA metabolic process" semantic networks, excluding terms associated with the obtained PN-related semantic groups.

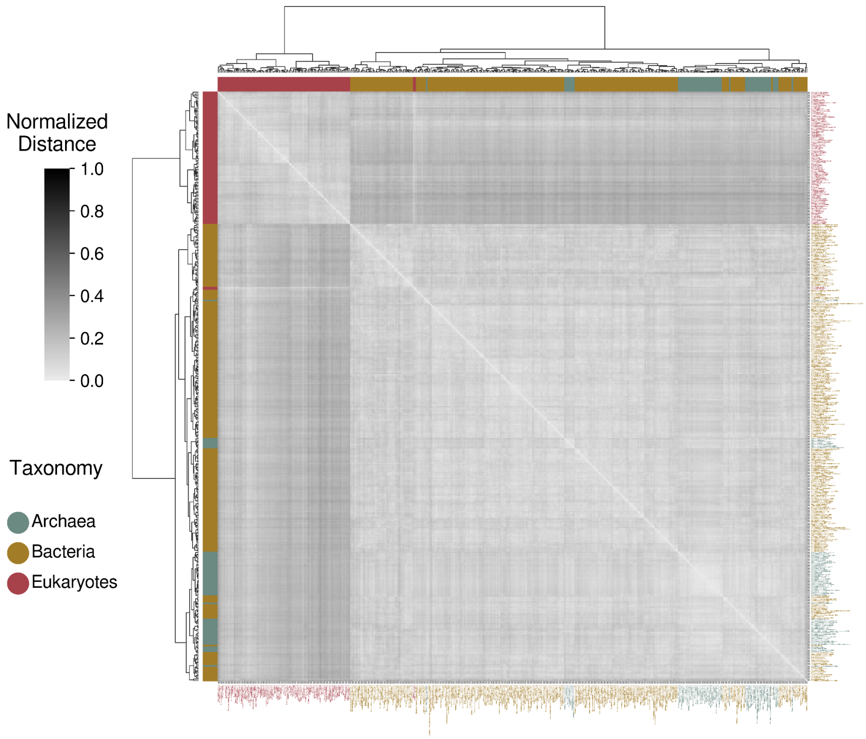


**Figure S2.30A**: Phylogenetic clustergram derived from the comparison of "ribonucleotide biosynthetic process" semantic networks.

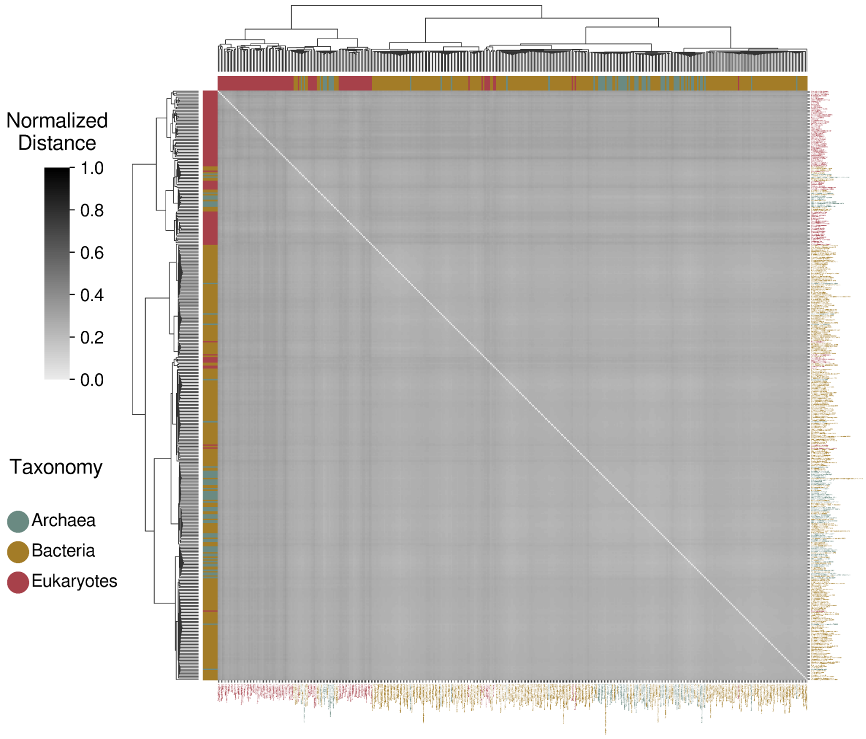


**Figure S2.30B**: Phylogenetic clustergram derived from the comparison of "ribonucleotide biosynthetic process" semantic networks, excluding terms associated with the obtained PN-related semantic groups.

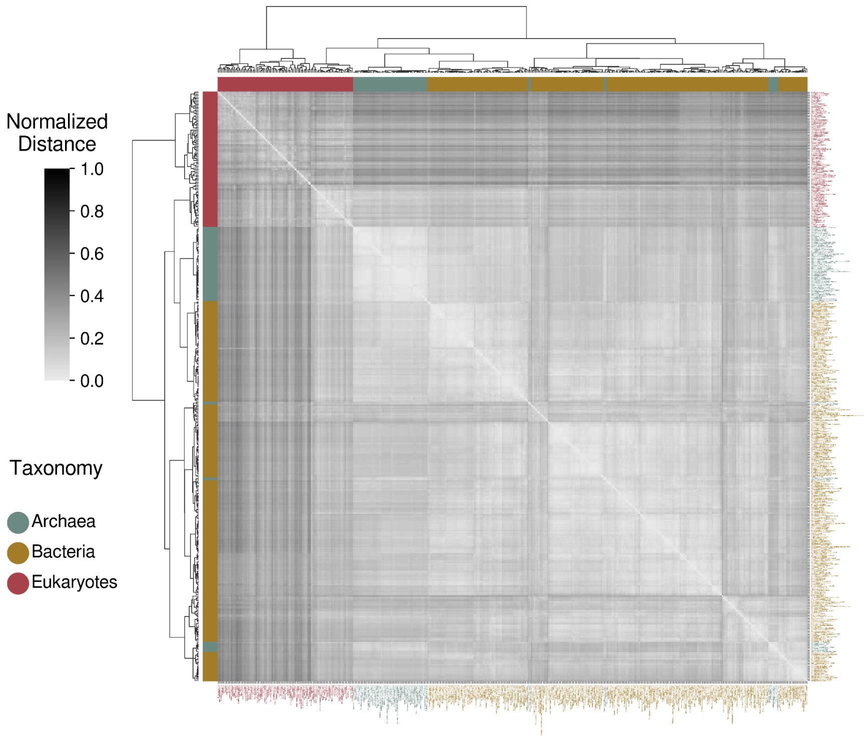


**Figure S2.31A**: Phylogenetic clustergram derived from the comparison of "tRNA processing" semantic networks.

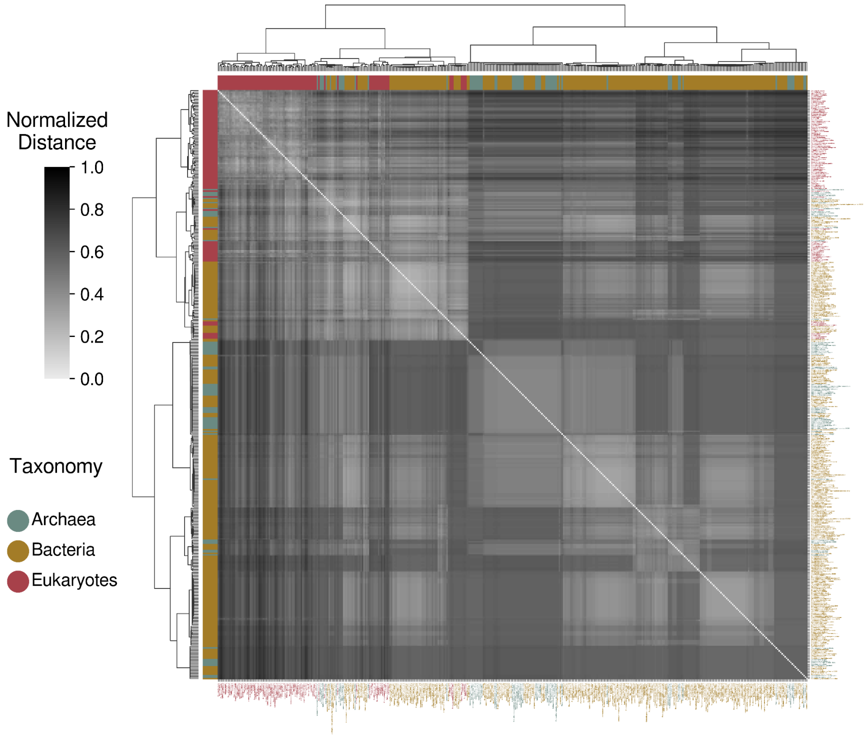


**Figure S2.31B**: Phylogenetic clustergram derived from the comparison of "tRNA processing" semantic networks, excluding terms associated with the obtained PN-related semantic groups.
